# Non-invasive measurement of mRNA decay reveals translation initiation as the major determinant of mRNA stability

**DOI:** 10.1101/214775

**Authors:** Leon Y Chan, Christopher F Mugler, Stephanie Heinrich, Pascal Vallotton, Karsten Weis

**Affiliations:** Department of Molecular and Cell Biology, University of California, Berkeley, Berkeley, United States; Department of Biochemistry, ETH Zurich, Zurich, Switzerland

## Abstract

The cytoplasmic abundance of mRNAs is strictly controlled through a balance of production and degradation. Whereas the control of mRNA synthesis through transcription has been well characterized, less is known about the regulation of mRNA turnover, and a consensus model explaining the wide variations in mRNA decay rates remains elusive. Here, we combine non-invasive transcriptome-wide mRNA production and stability measurements with selective and acute perturbations to demonstrate that mRNA degradation is tightly coupled to the regulation of translation, and that a competition between translation initiation and mRNA decay -but not codon optimality or elongation- is the major determinant of mRNA stability in yeast. Our refined measurements also reveal a remarkably dynamic transcriptome with an average mRNA half-life of only 4.8 minutes - much shorter than previously thought. Furthermore, global mRNA destabilization by inhibition of translation initiation induces a dose-dependent formation of processing bodies in which mRNAs can decay over time.

## Intro

Gene expression is responsible for the production of the macromolecular machinery required for life. It is thus the central process that drives all cellular processes and the amounts and modification states of the mRNA and protein gene products are what ultimately determine the identity, function and fate of a given cell. The abundances of both mRNAs and proteins are in turn determined kinetically by balancing both synthetic and degradative processes. At the mRNA level, we have a detailed understanding of both how mRNAs are made and how the individual steps of transcription, splicing and maturation are regulated. However, less is known about the regulation of mRNA decay and whereas individual steps of mRNA degradation have been determined, the question of what determines the stability of mRNAs across the transcriptome remains largely unanswered.

Bulk mRNA degradation was shown to be initiated by the removal of the polyA tail [1, 2]. This triggers degradation through one of two pathways. mRNAs can either be degraded from the 3’ end by the exosome complex of 3’ to 5’ exonucleases or -what is thought to be more common in yeast-deadenylation is followed by removal of the 5’-methylguanosine cap by the decapping complex [3, 4]. Removal of the cap structure is then followed by exonucleolytic digestion from the 5’ end of the mRNA by the cytoplasmic 5’ to 3’ exonuclease, Xrn1. While these pathways of mRNA degradation are well elucidated, their upstream regulators remain less clear and it is not well understood how the decision is made whether an mRNA continues to be translated or enters the decay pathway.

Factors ranging from polyA tail length to mRNA structure have been proposed to affect global transcript stability but many models have been centered on how the process of translation regulates transcript lifetime. Two alternative models have been put forth to explain how mRNA decay is linked to translation (Figure 3A). The first model originates from the observation that mRNA stability significantly correlates with codon usage. It was proposed that slowly elongating ribosomes at suboptimal codons signal to the decay machinery to target the bound mRNAs for destruction. Therefore, this stalled ribosome-triggered decay model centers on the process of translation elongation [5, 6]. The second model arises from the observations that translation and decay are inversely related and posits that bound translation factors protect an mRNA from decay. Such a translation factor-protection model predicts that translation initiation, either directly or indirectly, competes with the RNA decay machinery. In the latter model, the stability of a given transcript would be determined by a competition between the eIF4F initiation complex and the decapping complex for the 5’ methylguanosine cap, and/or by ribosomes sterically blocking decay factors from the mRNA [7–9]. Both of these models have supporting experimental evidence and are also not mutually exclusive. However, the available experimental evidence for each of these models has mainly been gathered using specific reporter transcripts and methods to measure mRNA stability that can introduce unintended effects and thus might lead to non-physiological measurements of half-life as discussed below. Moreover, the perturbations that have been employed to probe the relationship between translation and decay have the potential for significant secondary effects. Thus, improved methods to both measure mRNA stability as well as perturbing core elements of the translation machinery are required to evaluate the existing models.

Classical measurements of mRNA stability for multiple transcripts in parallel have required global inhibition of transcription. This leads to two major complications. The first is that the global inhibition of transcription is a major perturbation to the cell and this has been shown to induce a general stress response [10]. This stress response is elicited regardless of the method of global transcription inhibition be it by pharmacological or genetic means. The second complication arises from the fact that the methods used to shutoff transcription have off-target effects themselves.

Temperature shock or the use of transcriptional inhibitors such as phenantroline and thiolutin have unintended effects on cell physiology independent of transcriptional inhibition [10, 11]. At the individual transcript measurement level, the use of transcriptional shutoff via the regulatable *GAL* promoter has also been widely used. However, shutoff of the *GAL* promoter requires an acute carbon source shift and it has been shown that several key factors in the decay pathway such as Pat1, Dhh1, Ccr4 and Xrn1 are regulated in a carbon source-dependent manner [12, 13].

In this study, we have sought to determine how mRNA half-life is controlled on a transcriptome-wide level and have taken a two-pronged approach to study the relationship between translation and decay. First, we refine a metabolic labeling-based assay to measure mRNA lifetimes in a transcriptome-wide and non-invasive manner. Our new measurements reveal a much more dynamic transcriptome than previously measured by metabolic labeling with an average and median transcript half-life of only 4.8 and 3.6 minutes respectively. Next, we combine this measurement tool with both pharmacological and conditional genetic tools to directly perturb the processes of translation initiation and elongation. Our studies show that the competition between translation initiation and mRNA turnover determines the lifetime for an mRNA whereas slowing elongation globally leads to mRNA stabilization. At the cellular level, we find that the formation of processing bodies, sites where mRNAs are thought to be repressed and destroyed, is stimulated when translation initiation is attenuated suggesting that processing bodies form when mRNA clients are shunted into the degradation pathway.

## Results

### An improved non-invasive metabolic labeling protocol reveals that the yeast transcriptome is highly unstable

We and others previously demonstrated that thio-modified uracil nucleobases, such as 4-thio-uracil (4TU) are efficiently incorporated into nascent RNA [14–17]. Metabolic labeling with 4TU at appropriate concentrations has no adverse effect on cell proliferation and does not lead to major changes in gene expression [10, 15] (see extended technical supplement). Taking advantage of the thio-reacting group, labeled RNAs can then be biotinylated allowing for an affinity-based separation of thio-labeled mRNAs from unlabeled mRNAs in pulse-labeling experiments (Figure 1A). When this strategy is applied in a time resolved manner, it enables the direct measurement of mRNA decay kinetics in an unperturbed system.

**Figure 1:**
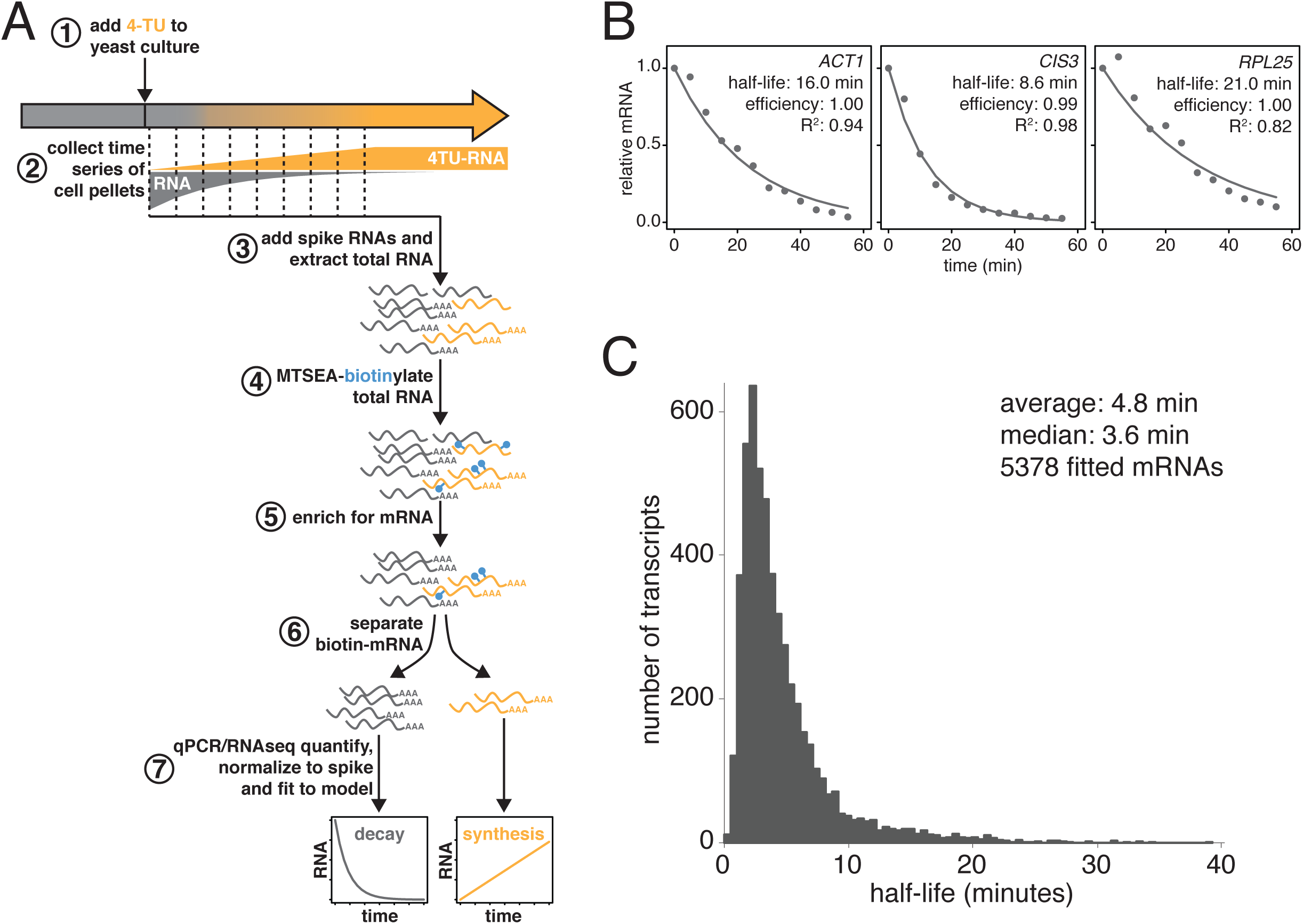
Improved mRNA stability measurements with metabolic labeling. (A) Experimental scheme to measure mRNA stability and production by metabolic labeling (B) Wild-type cells (KWY165) were subjected to the experiment described in (A) and decay rates for *ACT1*, *CIS3* and *RPL25* transcripts were determined by RT-qPCR. (C) mRNA samples in (B) were quantified by RNA-seq to determine mRNA stabilities across the transcriptome. Half-lives are plotted by frequency and each bin is 0.5 minutes wide.

We made three key modifications to our previously published protocol [14]. All modifications targeted the problem of inefficient subtraction of newly synthesized mRNAs which can result from inefficient chase of a metabolic label or low enrichment for labeled RNA during biotin-mRNA separation (see extended technical supplement for a detailed discussion). First, we optimized streptavidin-bead blocking and washing conditions that significantly reduced non-specific RNA bead binding (Figure 1A, step 6). Notably, the extent of non-specific bead binding differed between mRNAs indicating that non-specific binding can cause transcript-specific as well as global errors in decay rate determination. Second, we switched to the recently developed MTSEA-biotin which crosslinks biotin to a free thio group with far greater efficiency than the previously used HPDP-biotin [18] (Figure 1A, step 4). In combination, this improved efficiency allowed us to employ a 4-thiouracil(4TU)-chase labeling scheme in which rates of transcription and decay can be measured simultaneously (Figure 1A, step 1). Lastly, to correct for remaining inefficiencies in 4TU labeling, biotinylation and thio-RNA separation, we introduced an efficiency parameter into our decay model that allows more accurate rates to be extracted from the decay measurements (Figure 1A, step 7 and 1B).

Using this protocol, we measured half-life values for 5378 of the 6464 annotated transcripts in rapidly dividing budding yeast with a high agreement between biological replicates (Pearson correlation = 0.90) (Figure S1C). The analysis of the efficiency parameter showed that 92% of transcripts are labeled and separated with > 90% efficiency and 98% of transcripts are labeled and separated with > 80% efficiency indicating that our experimental optimizations are performing well (Figure S1D). Moreover, our data displayed a good fit to our modified exponential decay model with 84% of transcripts fitting with an R^2^ of > 0.95 (Figure S1E). Of the remaining transcripts, 209 fit the model poorly (R^2^ < 0.8) and an additional 877 could not be fit (Figure S1B). Most of these latter transcripts showed very low expression in our growth conditions and were thus excluded from the analysis.

The new measurements with our improved protocol revealed a much less stable transcriptome than previously reported, with average and median mRNA half-lives of 4.8 and 3.6 minutes respectively (Figure 1C). By summing the abundance of all mRNAs, we calculated the half-life of the bulk transcriptome to be 13.1 minutes (Figure S1A). Note that this value is higher than the 4.8 minute average value because it takes into account transcript abundance and many of the longest-lived transcripts are present in many copies within the mRNA pool. Importantly, this measurement agrees remarkably well with previous ^14^C-adenine pulse-labeling experiments, which are the least invasive measurements that have been performed to date, reporting a 11.5 minute half-life for the bulk polyA-RNA pool in the cell [19].

Consistent with the extensive protocol optimization, we found an overall poor correlation with our previously published dataset (Figure S1F). Nonetheless, our current measurements are consistent with the findings of Munchel et al. that long-lived (> 1 SD above the mean) transcripts are functionally enriched for translation factors and that ribosomal protein-encoding mRNAs specifically are long lived as a group with an average half-life of 15.5 minutes (Figures S1G and S1H). There is no significant functional enrichment in genes with exceptionally short (< 1 SD below the mean) mRNA half-lives. Our dataset does not agree well with the datasets derived from global transcriptional inhibition, which cluster with each other [11](Figure S1I). This is consistent with the findings of Sun et al. and Harigaya et al. that methods that rely on transcriptional inhibition all induce a global stress response that is elicited regardless of the method of transcriptional inhibition [10, 11]. Instead, our dataset clusters with the datasets of Cramer and Gresham that also employed non-invasive metabolic labeling although the transcriptome is much less stable by our measurements (Figure S1I) [10, 15, 20]. The overall distribution of half-lives for all fitted mRNAs (Figure 1C) is non-Gaussian stretching across more than an order of magnitude. The shortest half-lives are less than 1 minute whereas the most stable transcripts have half-lives of more than 30 minutes.

### Slowing translation elongation protects transcripts against degradation

To begin to identify factors that regulate this half-life diversity, we compared our decay dataset to other transcriptome-wide datasets of various mRNA measurements (Figure 2). Our decay data clustered with transcript abundance, metrics of codon usage (normalized translational efficiency (nTE) and codon adaptation index (CAI)), as well as translational efficiency measured by ribosome footprinting [21, 22]. The positive relationship between abundance and half-life supports the notion that mRNA levels are not only primarily dictated by the rate of synthesis, but that differential mRNA stability contributes to the regulation of transcript abundance as well. Interestingly, mRNA half-life was negatively correlated with polyA-tail length consistent with prior observations [23].

**Figure 2:**
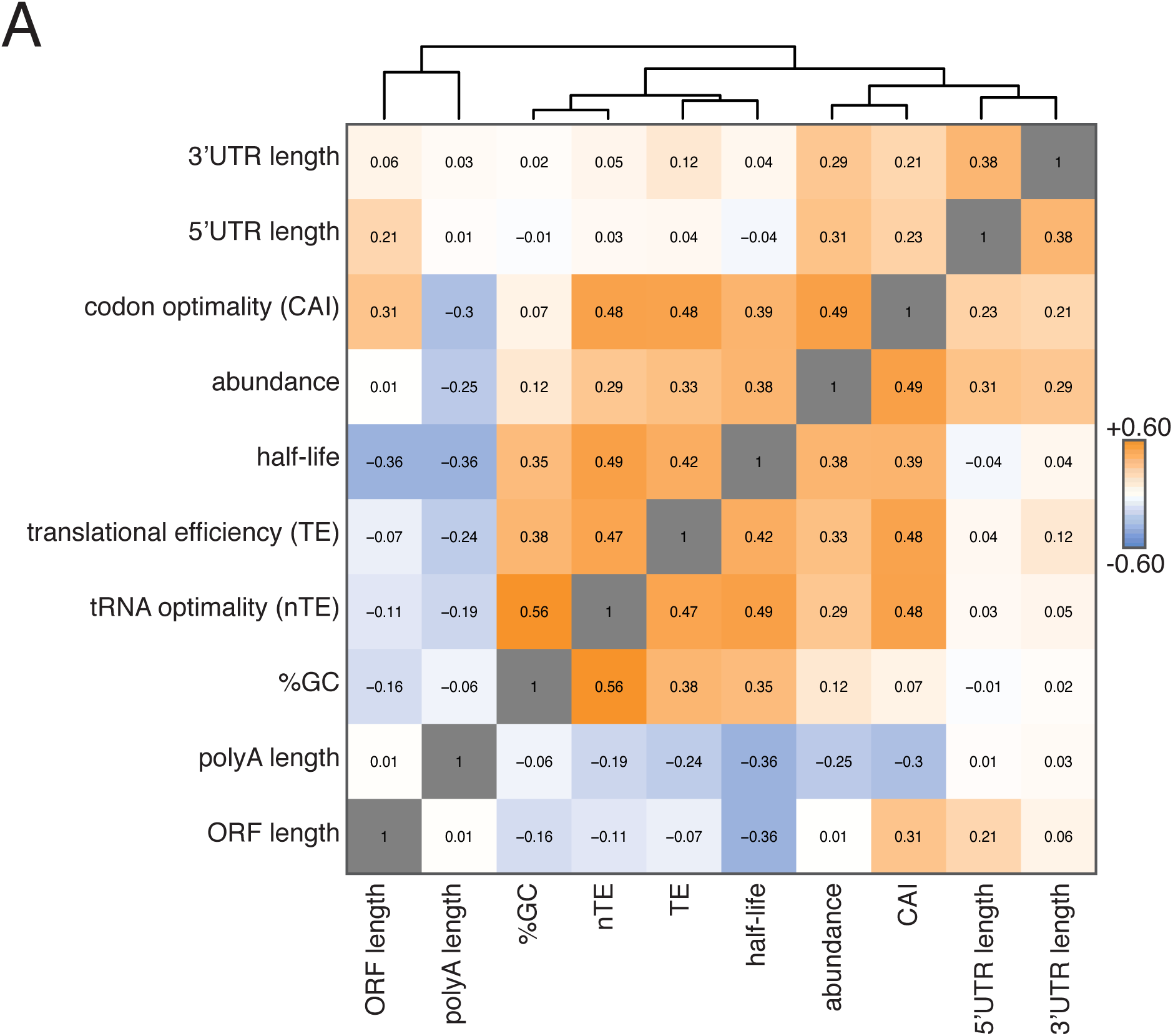
Correlation of mRNA features. (A) Spearman rank correlation coefficients were computed for pairs of mRNA parameters of stability (half-life), translation efficiency (TE), polyA tail length, codon optimality (CAI), tRNA optimality (nTE), abundance, UTR lengths, GC content and ORF length and plotted as a heatmap. Datasets were hierarchically clustered based on Euclidian distances. Orange represents positive correlation and blue represents negative correlation. Correlations between identical datasets are colored in gray. See supplemental table 1 for sources of genome wide data.

Our correlation analyses support prior work pointing to protein translation efficiency as a critical determinant of mRNA half-life. The aforementioned stalled ribosome-triggered decay and translation factor-protection models attempt to explain the positive correlations between mRNA half-life and codon usage and mRNA half-life and translation efficiency respectively (Figure 3A). These two models make clear and opposing predictions for how perturbing the processes of translation elongation or initiation impacts transcript stability. The stalled ribosome-triggered decay model predicts that mRNAs are destabilized upon slowing elongation whereas the translation factor-protection model predicts the opposite since slowly elongating ribosomes would accumulate on a given transcript and thus provide greater steric exclusion of decay factors. In contrast, when translation initiation rates are attenuated, the stalled ribosome-triggered decay model predicts that transcripts would either have the same stability or possibly even increased stability as once the bound ribosomes complete translation, the naked mRNA would be freed from decay-triggering ribosomes. The translation factor-protection model again predicts the opposite outcome: decreasing the rate at which translation is initiated leaves the 5’ cap more exposed to the decapping machinery and fewer loaded ribosomes allows the decay factors greater access to the transcript culminating in an overall decrease in transcript stability.

**Figure 3:**
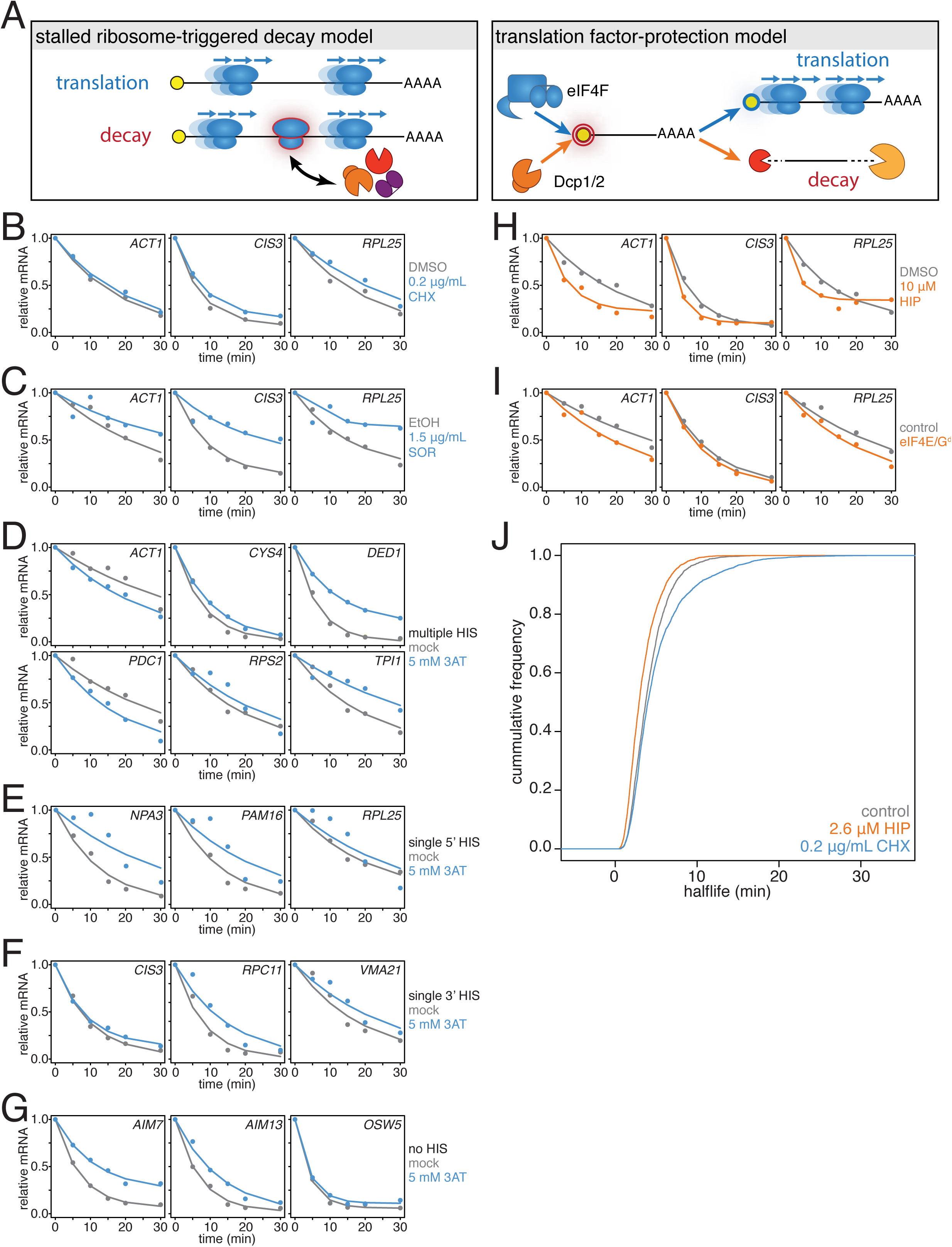
mRNAs are stabilized by slowly elongating ribosomes and destabilized when translation initiation is inhibited. (A) Cartoon depictions of the stalled ribosome-triggered decay and translation factor-protection model. (B) Wild-type cells (KWY165) were subjected to mRNA stability profiling immediately after addition of 0.1% DMSO or 0.2 μg/mL cycloheximide in 0.1% DMSO. Data on *ACT1*, *CIS3* and *RPL25* mRNAs were collected and plotted. (C) Wild-type cells (KWY165) were subjected to mRNA stability profiling 33 minutes after addition of 0.1% ethanol or 1.5 μg/mL sordarin in 0.1% ethanol (note that this is the timepoint when a growth defect is manifested, see figure S2C). Data were collected and plotted as in figure 3B. (D-G) *HIS3 gcn2∆* cells (KWY7337) were subjected to mRNA stability profiling immediately after non-addition (mock) or addition of 5 mM 3AT. Data were collected and plotted as in figure 3B. (H) Wild-type cells (KWY165) were subjected to mRNA stability profiling immediately after addition of 0.1% DMSO or 10 μM hippuristanol. Data were collected and plotted as in Figure 3B. (I) *pGPD1-LexA-EBD-B112 CDC33-3V5-IAA7* pRS425 cells (KWY7336: control) and *pGPD1-LexA-EBD-B112 CDC33-3V5-IAA7 pGPD1-OsTIR1* pRS425-*p4xLexOcyc1-CDC33*^*∆CAP*^ cells (KWY7334: eIF4E/G^down^) were grown in CSM-LEU-0.5xURA pH5.5 media and subjected to mRNA stability profiling immediately after addition of 10 nM β-estradiol, 100 μM 3-indoleacetic acid and 4 μM IP6. Data were collected and plotted as in Figure 3B. (J) Wild-type cells (KWY165) were subjected to global mRNA stability profiling immediately after addition of 0.1% DMSO (gray) or 2.6 μM hippuristanol (orange) or 0.2 μg/mL cycloheximide (blue). Cumulative frequencies of transcript half-life are plotted.

To begin to test these predictions, we directly perturbed the process of translation elongation and observed the effects on mRNA decay. Consistent with previous reports, we found that treating cells with a strong dose (50 μg/mL) of the elongation inhibitor cycloheximide resulted in stabilized *ACT1*, *CIS3* and *RPL25* transcripts (Figure S2A) [7]. A potentially problematic aspect of these experiments is that high doses of cycloheximide completely halt translation thus shutting down myriad cellular processes, which in turn could lead to indirect effects. To address this issue, we titrated the amount of cycloheximide to a sub-lethal concentration (0.2 μg/mL) and assayed the effect of this level of elongation inhibition on transcript stability (Figure S3H). Even with this low concentration of cycloheximide, the *ACT1*, *CIS3* and *RPL25* transcripts remained stabilized (Figure 3B). The effects of cycloheximide on mRNA turnover were specific to elongation inhibition as mRNA half-lives were unchanged in a cycloheximide resistant mutant (*rpl28-Q38K*) (Figure S2B). We next tested an alternative translation elongation inhibitor, sordarin, which blocks the function of eukaryotic elongation factor 2 [24]. When cells were treated with a sub-lethal dose of sordarin, we again observed a stabilizing effect (Figure 3C and S2C). These stabilization effects were not due to an inability to incorporate 4TU into newly made transcripts as mRNA synthesis rates were not reduced upon treatment with either cycloheximide or sordarin (Figures S2D and S2E). We conclude that inhibiting translation elongation stabilizes mRNAs.

While these results demonstrate that a stalled ribosome *per se* is not sufficient to induce decay, we could not exclude that cycloheximide or sordarin treatment might only poorly imitate slowed ribosomes on non-optimal codons since the acceptor-site of the ribosome remains occupied when these drugs are employed [25]. To best mimic a non-optimal codon where the acceptor-site would be unoccupied, we treated cells with a sub-lethal dose (5 mM) of 3-amino-1,2,4-triazole (3AT), which results in histidine starvation thus lowering the concentration of histidyl-tRNAs (Figure S2F) [26]. Indeed 3AT has previously been shown to stall ribosomes at histidine codons [27]. Histidine starvation also affects translation initiation by phosphorylation of eukaryotic initiation factor 2α via the Gcn2 kinase [28]. In order to examine the effect on translation elongation by 3AT in isolation, all 3AT experiments were thus performed in *gcn2∆* mutant cells. We examined a panel of 15 mRNAs with diverse spacing and position of histidine codons and found 13 of them to be stabilized whereas only 2 displayed increased decay kinetics upon 3AT treatment (Figure 3D-G). This overall stabilization effect could not be explained by poor 4TU uptake as mRNA synthesis rates were not reduced upon 3AT treatment (Figure S2G). Interestingly, transcripts lacking histidine codons were also stabilized, which is consistent with the observation that 3AT limits glycine availability in addition to the depletion of histidine [29]. Altogether, we conclude that inhibiting translation elongation by three different methods led to an overall stabilization of mRNAs. These observations are not consistent with the predictions of the stalled ribosome-triggered decay model but instead support a translation factor-protection model.

### Inhibition of translation initiation destabilizes individual transcripts

We next studied the effects of inhibiting translation initiation on mRNA decay. We first made use of hippuristanol, an inhibitor of eukaryotic initiation factor 4A (eIF4A) [30]. We observed that *ACT1*, *CIS3* and *RPL25* mRNAs all decayed with faster kinetics when eIF4A was inhibited (Figure 3H). To test the specificity of hippuristanol, we attempted to generate hippuristanol-resistant alleles of the eIF4A encoding genes, *TIF1* and *TIF2,* but these mutations (V326I, Q327G and G351T) led to severe cell sickness (data not shown) [31]. To exclude any potential indirect effects of hippuristanol, we sought alternative means to inhibit translation initiation. Overexpression of a 5’cap-binding mutant of eIF4E (*cdc33-W104F-E105A* henceforth *cdc33^∆CAP^*) using a β-estradiol inducible promoter caused a subtle inhibition of growth [32, 33]. This defect was fully suppressed by introducing in cis the ∆1-35 (henceforth *cdc33^∆G^*) mutation that abolishes eIF4G binding (Figures S3A and S3B) [34] indicating that overexpression of *cdc33*^*∆cap*^ leads to a dominant-negative loss of eIF4G function likely through a sequestration mechanism (Figure S3C). In addition, we placed eIF4E under control of an auxin-inducible degron system (*CDC33-3V5-IAA7*) [35]. This approach alone led to a mild growth defect upon the addition of auxin presumably because eIF4E could not be fully depleted (Figures S3D - F). However, when these two strategies were combined to simultaneously downregulate eIF4E and eIF4G function, we observed a strong synthetic growth defect (Figure S3G). This system then enabled us to acutely inhibit initiation in a manner orthogonal to hippuristanol and evaluate the resulting effects on mRNA decay. As with hippuristanol treated cells, we found that *ACT1*, *CIS3* and *RPL25* transcripts are all destabilized when translation initiation is slowed (Figure 3I). Based on the results of two independent experimental approaches we conclude that inhibiting translation initiation leads to accelerated mRNA decay.

### Translation elongation and initiation globally affect mRNA half-lives

To test the generality of our findings, we also performed transcriptome-wide mRNA stability profiling of cells treated with either cycloheximide or hippuristanol. To allow for a meaningful comparison, we used hippuristanol at a sub-lethal concentration that confers a near identical growth defect as our sub-lethal concentration of cycloheximide (Figure S3H). In support of our observations with individual mRNAs, cycloheximide induced a global stabilization of mRNAs (p = 6.298e-106 two-sided Wilcoxon paired test) whereas hippuristanol treatment led to shorter mRNA half-lives (p = 1.864e-260 two-sided Wilcoxon paired test) (Figure 3J). Importantly, the Spearman rank correlation coefficient between these datasets was high (Rsp(DMSO:HIP) = 0.81 and Rsp(DMSO:CHX) = 0.79). This suggests that these drugs did not result in a reordering of the stability profile of the transcriptome or differentially affect specific classes of mRNAs. Instead, this indicates that the drugs generally shifted the profile towards more (cycloheximide) or less (hippuristanol) stable. We conclude that slowing initiation accelerates mRNA turnover while inhibiting elongation slows mRNA turnover and that on a transcriptome-wide level, the efficiency of initiation either directly through 5’-cap competition or indirectly through ribosome protection is a major determinant of transcript stability.

### Inhibition of translation initiation induces processing bodies

What are the consequences of these perturbations to translation and their effect on mRNA decay at the cellular level? Inhibition of elongation with cycloheximide was previously shown to inhibit the formation of processing bodies (PBs), which are thought to be sites of transcript repression and decay [36–38]. To test the effects of inhibiting translation initiation on PB formation, we treated cells expressing Dhh1-GFP and Dcp2-mCherry markers of PBs with a range of hippuristanol concentrations. Interestingly, hippuristanol induced PB formation in a concentration dependent manner: at high doses (10 – 40 μM), rapid and robust PB formation could be observed; at an intermediate dose (5 μM), PBs formed over time and at a low dose (2.5 μM), PBs could not be detected (Figure 4A and 4B). Since hippuristanol generates client mRNAs for the decay machinery through its inhibition of initiation, this dosage effect suggests that PB formation is directly dependent on the number of mRNA substrates available for degradation and that microscopic PBs can only be detected when a certain threshold of decay targets is reached. Consistent with such a model, we observed the rapid relocalization of three distinct mRNAs, *GFA1*, *PGK1* and *FBA1,* to PBs upon hippuristanol-induced PB formation (Figure 4D). Unlike in mammalian cell culture systems, hippuristanol does not trigger the formation of stress granules in yeast (Figure S4A) but as with other PBs, the formation of hippuristanol-induced Dhh1- and Dcp2-containing foci requires the RNA and ATP binding activities of Dhh1 (Figures S4B and S4C) [38, 39]. An alternate explanation for these hippuristanol-induced PBs is that the perturbation of translation alone may result in cellular stress and PB formation. However, co-treatment of hippuristanol-treated cells with either cycloheximide or sordarin suppressed PB formation, suggesting that the increased number of ribosome-unbound mRNA clients available for degradation, rather than crippled translation, was causative for PB formation (Figure 4A and 4C).

**Figure 4:**
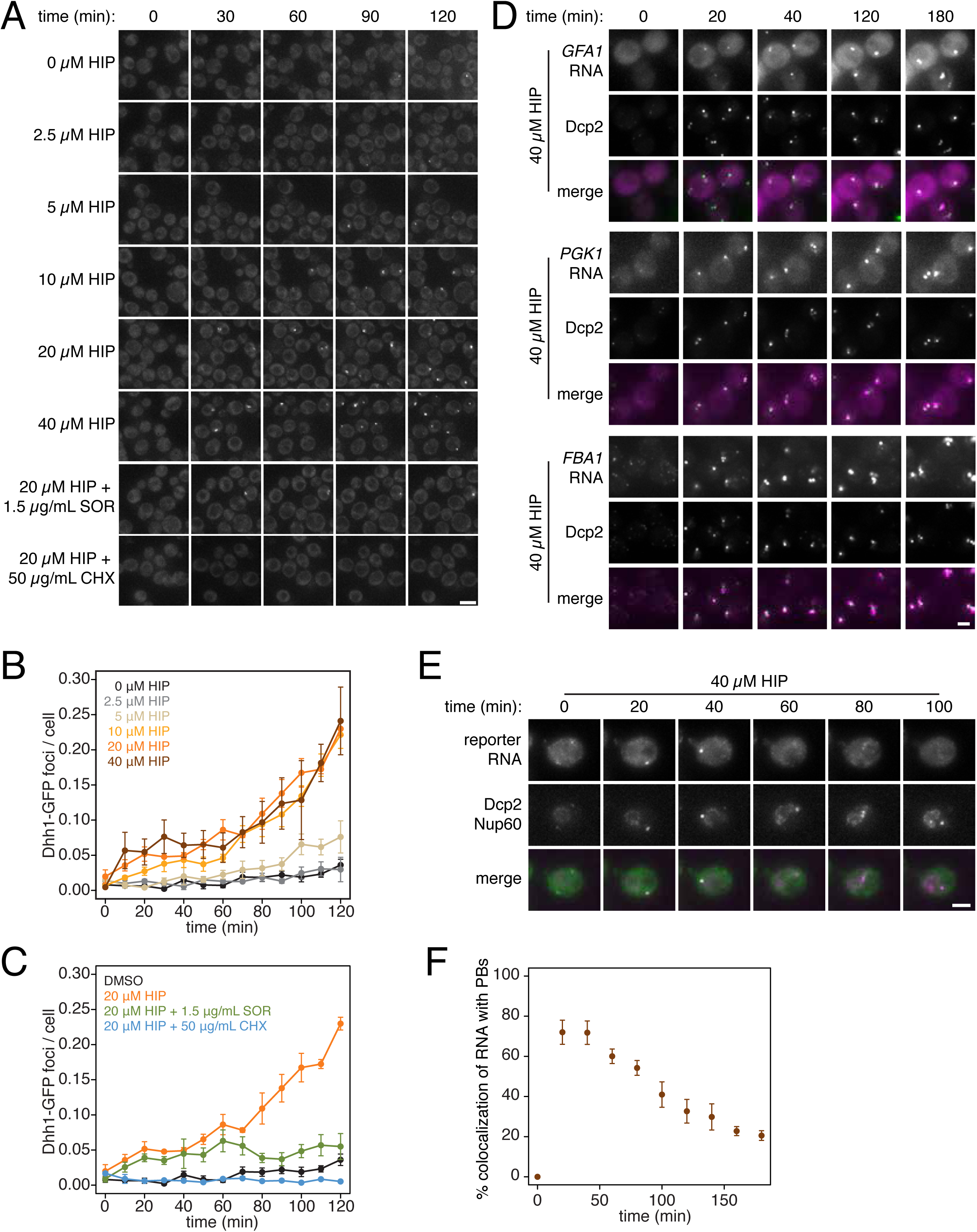
PB formation is stimulated by inhibiting translation initiation and blocked when translation elongation is inhibited. (A) Dhh1-GFP, Dcp2-mCherry expressing cells (KWY5948) were grown to exponential phase and then treated with 0.1% DMSO, the indicated concentration of hippuristanol or co-treated with the indicated concentration of hippuristanol and either sordarin or cycloheximide. Images were aquired every 5 minutes using a wide-field microscope and the images were deconvolved. Shown are maximum projections of 8 z-stacks at a distance of 0.4 μm apart. Scale bar: 5 μm. (B-C) Number of Dhh1-GFP foci per cell from experiment in (A) was counted using Diatrack 2.5 particle tracking software. Error bars represent SEM (n= 3 biological replicates, > 300 cells counted per experiment). (D) Dcp2-GFP, PP7CP-mKate2 expressing cells carrying PP7sl tagged copies of *GFA1* (KWY7246), *PGK1* (KWY6963) or *FBA1* (KWY7245) were treated with 40 μM hippuristanol and immediately imaged. Images where acquired every 20 minutes using a wide-field microscope. Shown are maximum projections of 8 z-stacks at a distance of 0.5 μm apart. Scale bar: 2 μm. (E) Dcp2-mCherry, Nup60-3xmKate2, PP7CP-GFP expressing cells carrying a synthetic 3xGST-24xPP7sl under β-estradiol inducible control (KWY7227) were grown to mid-exponential phase, treated with 400 nM β-estradiol for 40 minutes and then transferred to media lacking β-estradiol and containing 40 μM hippuristanol and immediately imaged (see supplemental Figure 4D for the no hippuristanol control). Images were acquired every 20 minutes using a wild-field microscope. Shown are maximum projections of 8 z-stacks at a distance of 0.5 μm apart. Scale bar: 5 μm. For DMSO control images, see figure S5D. (F) Images acquired in (E) were quantified for the colocalization of PP7CP-GFP foci with Dcp2-mCherry foci using FIJI software. Error bars represent SEM (n = 4 biological replicates, > 120 PBs counted per timepoint).

Recent evidence has supported the notion that mRNAs can be degraded in PBs [38, 40]. To examine whether the PBs that form upon addition of hippuristanol can be sites of mRNA degradation, we placed a model transcript containing slowly decaying PP7 stem loops (PP7sl) under control of a β-estradiol inducible promoter [40]. We pulsed cells with this transcript by treating the cell for 40 min with β-estradiol, washed out the inducer, immediately added 40 μM hippuristanol and then observed the localization of the PP7 stem loops over time. As observed for endogenous mRNAs, we found that the PP7sl-containing transcript rapidly localized to PBs (Figure 4E). Moreover, we observed that the PP7-mRNA signal decayed over time within the PB (Figure 4E and 4F). This suggests that mRNAs localize to PBs when initiation is inhibited and that these mRNAs can decay over time within the PB. In combination with our metabolic labeling studies, we further conclude that inhibiting translation initiation leads to global mRNA destabilization which in turn causes the formation of PBs. In the presence of agents that inhibit translation elongation, mRNAs become stabilized reducing the flux of new client mRNAs into the degradation pathway, which in turn suppresses the formation of PBs.

### Discussion

In this work, we have refined an assay to measure the kinetics of mRNA synthesis and decay based on 4TU metabolic labeling. This approach and similar approaches supersede the traditional methods of transcriptional inhibition as they enable quantitative and global measurements of mRNA kinetics in physiologically unperturbed cells. We used this approach to address the important question of how the process of translation affects transcript stability. Importantly, all of the measurements and experimental perturbations employed here relied on minimally invasive and rapidly inducible methods. Moreover, the drugs we used have specific molecular targets and the genetic inhibitions of eIF4G and eIF4E are induced by hormones from orthologous systems, which have minimal off-target effects.

Taken together, these approaches have enabled us to identify translation initiation as the central hub in globally regulating mRNA stability. While no direct measurement of translation initiation rates at the transcriptome-wide level has been reported, rates estimated by using a kinetic model of ribosome flux [41] are in agreement with our conclusion and the dataset of estimated rates strongly correlates and clusters most closely with transcript half-life (Figures S5A and S5B). While we cannot exclude that suboptimal codons enhance the decay of specific transcripts, our results do not support the hypothesis that stalled ribosomes are the primary contributor to cellular mRNA decay on a transcriptome-wide level.

Nevertheless, we did find a positive correlation between codon optimality and transcript stability as previously reported [5]. However, similar positive correlations can also be identified with general translation efficiency and transcript abundance. We therefore consider it likely that these properties have undergone similar selection pressure and might have co-evolved with transcript stability to ensure optimal gene expression of particular highly abundant transcripts.

It has been previously proposed that deadenylation is the rate limiting step of mRNA decay [42]. If this was the case and the rate of deadenylation for each transcript was constant, one would expect that the length of the polyA tail would directly determine the stability of the associated transcript. However, rather than positively correlating with half-life, we find that polyA tail length negatively correlates with transcript stability consistent with prior results [23]. Despite this inverse relationship, it is important to note the negative effects of polyA-binding protein on transcript decapping and thus the role of deadenylation and the length of the polyA tail in controlling transcript stability are likely more nuanced than a simple rate-limiting model would imply [43, 44]. Moreover, it will be important to examine not only a snapshot of the steady state polyA tail length but to determine the kinetics of polyA tail shortening to understand if and how the rate of deadenylation contributes to overall transcript stability.

Our work also suggests that a sudden increase of decay clients leads to PB formation once a critical threshold is reached. This is consistent with previous studies showing that mRNA is required for PB formation and further implies that mRNA can be limiting for PB formation when translation is rapidly downregulated as is the case during cellular stress. Furthermore, as mRNA decay and translation are opposing fates for an mRNA and are competing processes in the cell, it might also be the case that the cell physically compartmentalizes these processes away from one another by use of a liquid-liquid phase transition droplet such as a P-body. A remaining open question is whether PBs form because the decay machinery is overburdened and decay intermediates accumulate or whether decay substrates are delivered to PBs in order to accelerate their decay. Inducing PBs by hippuristanol treatment will be an excellent approach to dissect these possibilities as this mode of PB formation bypasses the typical stresses associated with PB formation such as nutrient starvation that might have more pervasive and confounding effects on mRNA decay itself. The role of PBs in mRNA turnover has remained unclear and controversial. Our data support the notion that mRNAs can be degraded in PBs but whether this is the primary function of PBs remains an open question. Moreover, these data add to a growing picture of mRNA turnover occurring in many intracellular locations including on the polysome, in translationally silenced mRNPs as well as in larger PBs [45–47]. Going forward, it will be important to measure the contribution of each of these decay sites to the overall capacity of the cell to destroy mRNAs, and to elucidate the cellular function(s) of PBs.

A revelation from this work is the overall short half-life of the transcriptome, only 4.8 minutes or a mean lifetime of 6.9 minutes. This value is 3 times faster than was previously measured by metabolic labeling and up to 26 times faster than what was measured by transcriptional inhibition. Despite these very short half-lives, with an estimated average translation initiation rate of 0.12 s^−1^, this implies that the average transcript can still code for about 50 polypeptides before it is destroyed [41]. This overall instability of the transcriptome argues against the need for regulated mRNA decay for the bulk of transcripts in the cell. That being said, there is a class of long lived transcripts that we and others have found to be enriched for translation factors and ribosomal protein encoding mRNAs, and there is indeed mounting evidence that these transcripts can have dramatically differing stabilities depending on the state of the cell [48, 49]. It is also important to note that our measurements were made in rapidly dividing yeast cells, and it remains to be examined whether the determinants of mRNA stability as well as the degree of regulated turnover could shift as cells are exposed to stresses or undergo differentiation programs. Our non-invasive metabolic labeling approach can be applied in such contexts to determine how decay and synthesis work together to kinetically shape dynamic gene expression programs.

## Experimental Procedures

### Yeast Strains and Growth Conditions

All strains are derivatives of W303 (KWY165) with the following exceptions: KWY7227 and KWY7246 are derivatives of BY4741 (KWY1601). All strains are listed in Supplemental Table 2. *gcn2∆*, *CDC33-IAA7-3V5*, *FBA1-PP7sl*, *GFA1-PP7sl* and *PGK1-PP7sl* were generated by standard PCR based methods (Longtine). *RPL28(Q38K)* was generated by plating wild-type cells on 3 mg/mL cycloheximide plates, selecting for suppressors, backcrossing the suppressors at least three times and confirming the mutation by sequencing. *HIS3* was generated by PCR replacement of the *his3-11,15* allele using *HIS3* from pRS303. *leu2-3,112∆::CG-LEU2::pGPD1-OsTIR1*, *his3-11,15∆::CG-HIS3::pGPD1-OsTIR1*, *trp1-1∆::CG-TRP1::pGPD1-LexA-EBD-B112*, *his3-11,15∆::CG-HIS3::pGPD1-LexA-EBD-B112* and *SCO2::p4xLexOcyc1-3xGST-V5-24xPP7sl-tCYC1-NatNT2* were generated by transforming strains with plasmids pKW2830 (PmeI digested), pKW2874 (PmeI digested), pKW3908 (SwaI digested), pKW4073 (SwaI digested) and pKW4190 (NotI/AscI digested) respectively. Strains were grown in CSM-lowURA (7 g/L YNB, 2% dextrose, 20 mg/L adenine, 20 mg/L arginine, 20 mg/L histidine, 60 mg/L leucine, 30 mg/L lysine, 20 mg/L methionine, 50 mg/L phenylalanine, 200 mg/L threonine, 20 mg/L tryptophan, 30 mg/L tyrosine, 10 mg/L uracil) unless otherwise indicated. The following chemicals were obtained from the indicated sources: cycloheximide [Sigma], hippuristanol [a generous gift of Junichi Tanaka, University of the Ryukyus], β-estradiol [Sigma], sordarin [Sigma], 3-indoleacetic acid [Sigma], IP6 [Sigma], 4-thiouracil (4TU) [Arcos], 3-amino-1,2,4-triazole (3AT) [Sigma].

### Plasmid Construction

All plasmids are listed in Supplemental Table 3. Plasmid sequence files will be provided upon request. pNH604-*pGPD1-LexA-EBD-B112* (pKW3908) was constructed by standard restriction enzyme cloning using plasmid FRP880 as a PCR template for LexA-EBD-B112 (Stelling) and pNH603-*pGPD1-LexA-EBD-B112* (pKW4073) was derived from this plasmid. pNH603-*pGDP1-OsTIR1* (pKW2874) and pNH605-*pGPD1-OsTIR1* (pKW2830) were constructed by standard restriction enzyme cloning using pNHK53 as a PCR template for OsTIR1 (Nishimura et al 2009 Nat Methods). pFA6a-*IAA7-3V5-KanMx6* (pKW4325) was generated by Gibson assembly using a cDNA pool of aradopsis thalania as template for IAA7. Plasmids pRS425-*p4xLexOcyc1-CDC33*(± *∆G* ±*CAP*)-(±*3V5*) (pKW4326, pKW4327, pKW4328, pKW4329, pKW4330, pKW4331, pKW4332 and pKW4333) were generated by a combination of Gibson assembly and site-directed mutagenesis using FRP793 (stelling) as a PCR template for p4xLexOcyc1. Plasmid pRS313-*HR1_Chr2(SCO2)-p4xLexOcyc1-3xGST-V5-24xPP7sl-tCYC1-NatNT2-HR2_Chr2(SCO2)* (pKW4910) was constructed using standard restriction enzyme based cloning methods.

### 4TU metabolic labeling and RNA analysis

Cells were grown in CSM-lowURA overnight to post-diauxic stage (OD600 = 1 – 5) and then backdiluted in CSM-lowURA at OD600 = 0.1. Cells were grown for at least two doublings, backdiluted to OD600 = 0.4 and 1 mM 4TU was added from a 1 M 4TU stock in DMSO. Cells were colleced by filtration and immediately snap frozen in liquid nitrogen. Cell pellets were resuspended in 400 μL ice-cold TES buffer (10 mM TrisHCl pH7.5, 10 mM EDTA, 0.5% SDS) containing 5 ng 4TU-*srp1α*(Hs) -polyA spike RNA and 5 ng *rcc1*(Xl)-polyA spike RNA. 400 μL acid-saturated phenol was added and samples were continuously vortexed for 1 hr at 65°C. The aqueaous phase was then subjected to an additional phenol extraction followed by chlorofrom extraction and then isopropanol precipitated. Total RNA was pelleted, resuspended and 14 μg was biotinylated with MTSEA-biotin [Biotium] as previously described [1]. 10 μg of biotinylated total RNA was then subjected to oligo-dT bead [Life technologies] selection and used as input for the steptavidin bead selection. 25 μL MyOne streptavidin C1 Dynabeads [Life technologies] were washed with 25 μL each of 0.1 M NaOH (2x), 0.1 M NaCl (1x) and buffer3 (10 mM TrisHCl pH7.4, 10 mM EDTA, 1 M NaCl) (2x). The beads were then resuspended in 25 μL buffer3 and 2.5 μL 50x Denhardt’s reagent was added. Beads were then incubated with gentle agitation for 20 minutes, washed with 75 μL buffer3 (4x) and resuspended in 25 μL buffer3 with 2 μL 5 M NaCl added. Oligo-dT selected biotinylated RNAs were added to the blocked streptavidin beads and incubated with gentle agitation for 15 minutes. The flowthrough was collected and the beads were washed with 75 μL each buffer3 prewarmed to 65°C (1x), buffer4 (10 mM TrisHCl pH7.4, 1 mM EDTA, 1%SDS) (1x) and 10%buffer3 (2x). The washes were pooled with the flowthrough and 25 μL 5 M NaCl and 15 μg linear acrylamide [Ambion] were added prior to isopropanol precipitation. Biotinylated RNAs were eluted from the streptavidin beads first by a 5 minute incubation with gentle agitation in 5% β-mercaptoethanol and a subsequent 10 minute incubation at 65°C in 5% β-mercaptoethanol. Eluates were pooled and 7 μL 5 M NaCl and 15 μg lineaer acrylamide were added prior to isopropanol precipitation. RNAs were DNaseI [NEB] treated prior to downstream analysis. 4TU-*srp1α*(Hs)-polyA and *rcc1*(Xl)-polyA spike RNAs were generated as previously described using plasmids pKW1644 and pKW1643 respectively [2].

### RT-qPCR quantification of RNA abundance

2-50 ng of mRNA (depending on sample) was used for reverse transcription using Superscript II [Life technologies] with random hexamer primers. cDNA was quantified on a StepOnePlus Real-Time PCR system using a SYBR green PCR mix [ThermoFisher] with gene specific primers (Supplemental Table 4).

### High-throughput sequencing and RNAseq quantification

50-100 ng mRNA was used as input material to generate strand-specific sequencing libraries using the NEXTflex Rapid Directional Illumina RNA-Seq Library Prep Kit [BioO] according to the manufacturer’s instructions. The libraries were sequenced on an Illumina HiSeq 4000 sequencer by the QB3 Vincent J. Coates Genomics Sequencing Laboratory. Reads were mapped, aligned and quantified using the Tuxedo tools [3].

### Immunoblot analysis

For immunoblot analysis of Cdc33-3V5 and Cdc33-IAA7-3V5, cell were pelleted and resuspended in 5% ice-cold trichloroacetic acid for a minimum of 10 minutes. The acid was washed away with acetone and cell pellets were air-dried overnight. The pellet was resuspended in 100 μL lysis buffer (50 mM Tris-HCl pH 7.5, 1 mM EDTA, 2.75 mM DTT) and pulverized with glass beads in a beadmill [BioSpec] for 5 minutes. Sample buffer was added and the lysates were boiled for 5 minutes. V5 tagged proteins were detected with a mouse anti-V5 antibody [Life technologies] at a 1:2000 dilution. Hexokinase (Hxk1) was detected using a rabbit anti-hexokinase antibody [H2035, Stratech] at a 1:10,000 dilution. IRdye 680RD goat-anti-rabbit [LI-COR Biosciences] and IRdye 800 donkey-anti-mouse [LI-COR Biosciences] were used as secondary antibodies.

### Data processing and half-life determination

RNA measurements were first normalized by the abundance of the spike RNA (*rcc1*(Xl) for decay measurements and *srp1α*(Hs) for synthesis measurements) and then normalized to the t = 0 value. These data were then fit to the following decay model:

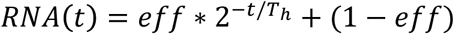

Where *eff* is a bulk efficiency term that accounts for the efficiency of 4TU labeling, biotin conjugation and separation of biotinylated RNAs from unbiotinylated RNAs and *T*_*h*_ is the half-life. A script was written in R to perform filtering, normalization and fitting of the decay data. For synthesis rate determination, data were fit to:

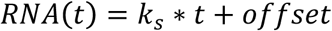

Where *k*_*s*_ is the synthesis rate and the *off set* is the y-intercept. Please see the extended technical supplement for more details.

### Growth determination

Cells were grown to mid-log phase, back diluted to OD 0.1-0.3 and growth at 30°C was monitored either manually by measuring absorbance at 600 nm or in a Tecan Infinite M1000 plate reader [Tecan].

### Wide-field fluorescence microscopy

Cells for imaging experiments were grown to exponential phase in CSM + 2% dextrose. Cells were then transferred onto Concanavalin A-treated 96-well glass bottom Corning plates (Corning, Corning, NY). For experiments described in Figures 4A-C and Supplemental Figure 4B-C cells were visualized at room temperature using the DeltaVision Elite Imaging System with softWoRx imaging software (GE Life Sciences, Marlborough, MA). The system was based on an Olympus 1X71 inverted microscope (Olympus, Japan), and cells were observed using a UPlanSApo 100 × 1.4 NA oil immersion objective. Single plane images were acquired using a DV Elite CMOS camera. For experiments described in Figures 4D-F and Supplemental Figure 4D, cells were visualized at room temperature using an inverted epi-fluorescence microscope (Nikon Ti) equipped with a Spectra X LED light source and a Hamamatsu Flash 4.0 sCMOS camera using a 100x Plan-Apo objective NA 1.4 and the NIS Elements software. Quantification of co-localization was performed on all planes of a 3D stack using the Colocalization Threshold tool in FIJI. Image processing for PB counting was performed using Diatrack 3.5 particle tracking software.

### Spinning disk confocal microscopy

Samples were grown in CMS + 2% dextrose to exponential phase and imaged using an Andor/Nikon Yokogawa spinning disk confocal microscope (Belfast, United Kingdom) with Metamorph Microscopy Automation & Image Analysis software (Molecular Devices, Sunnyvale, CA). The system was based on a NikonTE2000 inverted microscope and cells were observed using a PlanApo100 × 1.4 NA oil immersion objective and single plane images were captured using a Clara Interline CCD camera (Andor).

## Acknowledgements

We are grateful to Dr. Junichi Tanaka for the generous gift of hippuristanol and to Dr. Jörg Stelling for generously sharing plasmids. We would like to thank Dr. Kathy Collins, members of the Ünal, Brar and Weis labs for their critical reading of this manuscript and helpful discussions, and the Brar and Ünal labs for their geneous sharing of equipment and reagents. LYC is an HHMI Fellow of the Damon Runyon Cancer Research Foundation and is further supported by a grant from the Shurl and Kay Curci foundation. SH acknowledges support from an EMBO long-term fellowship (ALTF 290-2014, EMBOCOFUND2012, GA-2012-600394 to SH). This work was supported by NIH/NIGMS (R01GM058065 and R01GM101257 to KW) and the Swiss National Science Foundation (SNF 159731 to KW). This work used the Vincent J. Coates Genomics Sequencing Laboratory at UC Berkeley, supported by NIH S10 OD018174 Instrumentation Grant.

---

**Supplemental Table 1.**
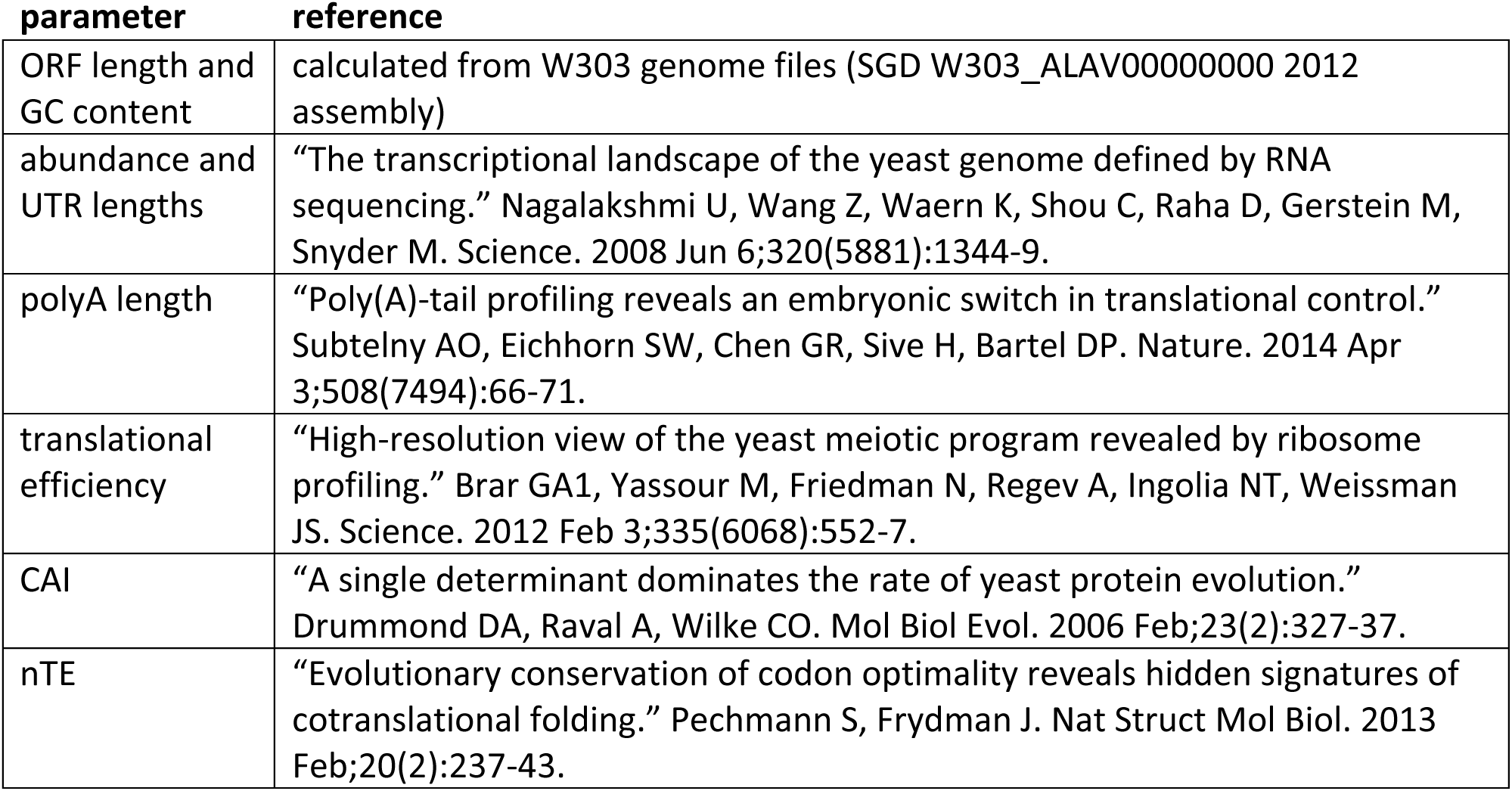

| **parameter** | **reference** |
| --- | --- |
| ORF length and GC content | calculated from W303 genome files (SGD W303_ALAV000000000 2012 assembly) |
| abundance and UTR lengths | "The transcriptional landscape of the yeast genome defined by RNA sequencing." Nagalakshmi U, Wang Z, Waern K, Shou C, Raha D, Gerstein M, Snyder M. Science. 2008 Jun 6;320(5881):1344-9. |
| polyA length | "Poly(A)-tail profiling reveals an embryonic switch in translational control." Subtelny AO, Eichhorn SW, Chen GR, Sive H, Bartel DP. Nature. 2014 Apr 3;508(7494):66-71. |
| translational efficiency | "High-resolution view of the yeast meiotic program revealed by ribosome profiling." Brar GA1, Yassour M, Friedman N, Regev A, Ingolia NT, Weissman JS. Science. 2012 Feb 3;335(6068):552-7. |
| CAI | "A single determinant dominates the rate of yeast protein evolution." Drummond DA, Raval A, Wilke CO. Mol Biol Evol. 2006 Feb;23(2):327-37. |
| nTE | "Evolutionary conservation of codon optimality reveals hidden signatures of cotranslational folding." Pechmann S, Frydman J. Nat Struct Mol Biol. 2013 Feb;20(2):237-43. |

**Supplemental Table 2.**
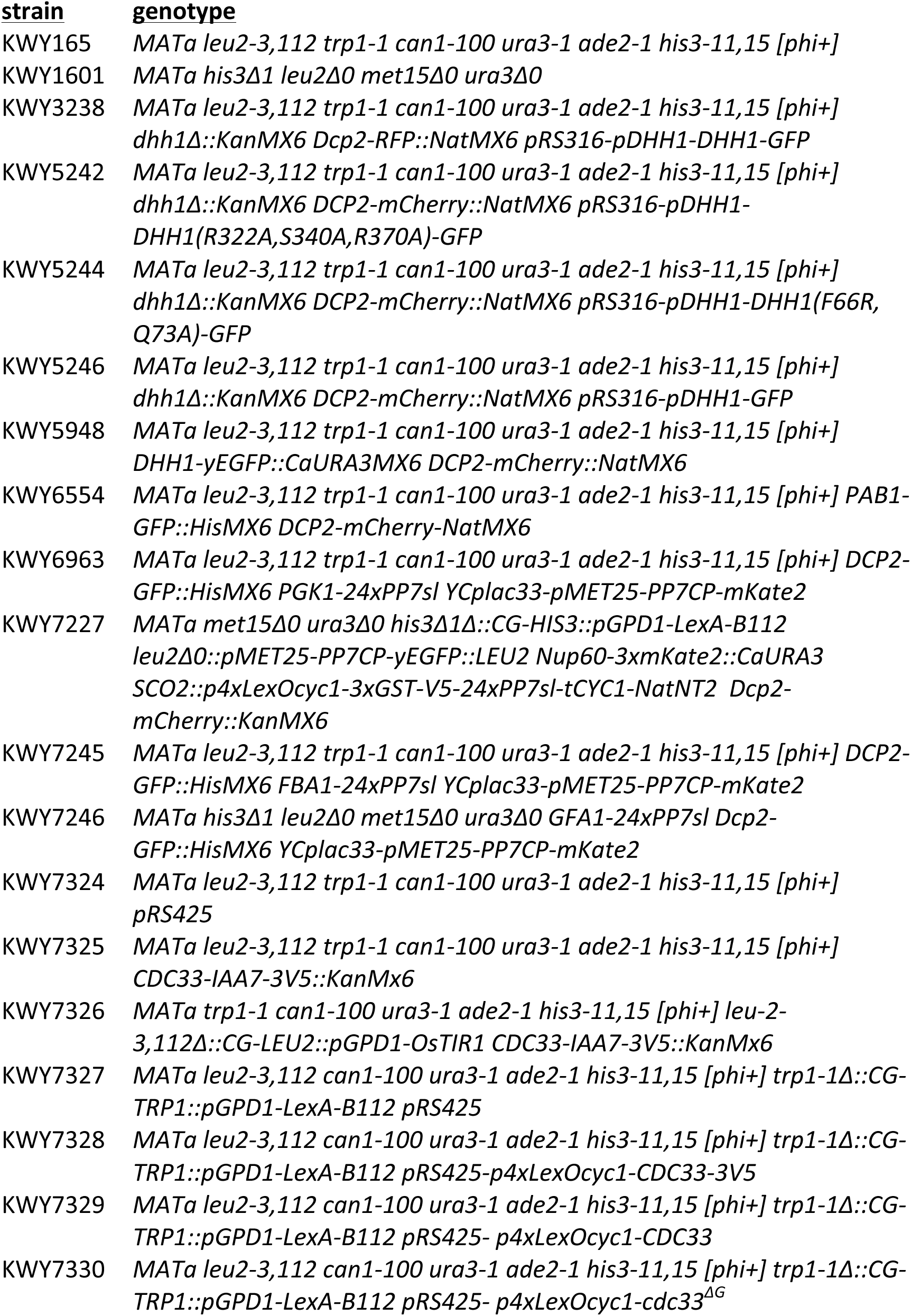
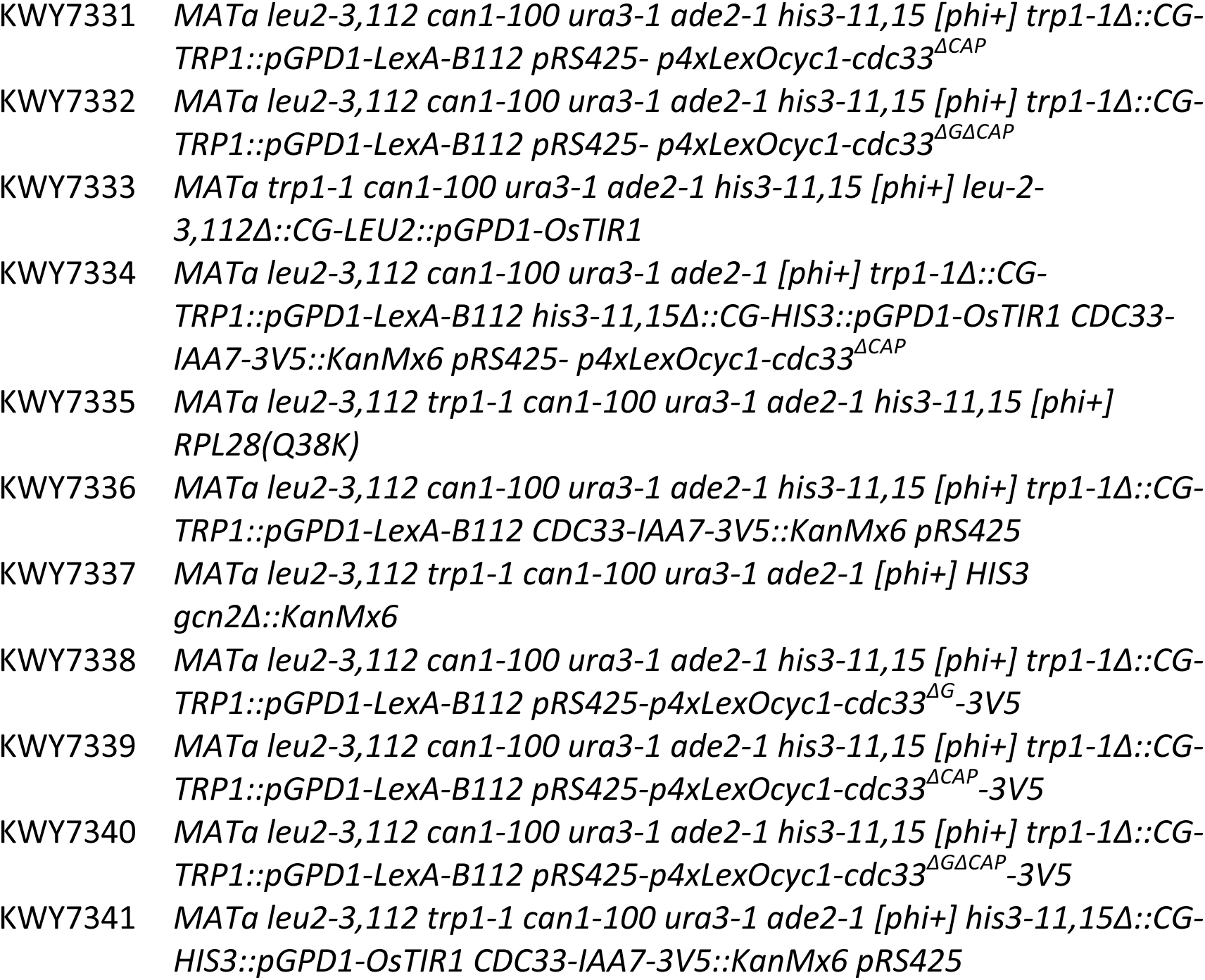

**Supplemental Table 3.**

|  |  |
| --- | --- |
| pKW1643 | pAS439-*rcc1* (Xl) |
| pKW1644 | pAS439-*srp1α* (Hs) |
| pKW2306 | pRS316-*pDHH1-GFP* |
| pKW2312 | pRS316-*pDHH1-DHH1-GFP* |
| pKW2550 | pFA6a-24xPP7sl-*tCYC1-KanMX6* |
| pKW2830 | pNH605-*pGPD1-OsTIR1* |
| pKW2867 | pRS316-*pDHH1-DHH1(R322A,S340A,R370A)-GFP* |
| pKW2874 | pNH603-*pGPD1-OsTIR1* |
| pKW3072 | pRS316-*pDHH1-DHH1(F66R, Q73A)-GFP* |
| pKW3616 | pRS305-*pMET25-PP7CP-yEGFP* |
| pKW3800 | YCplac33-*pMET25-PP7CP-mKate2* |
| pKW3908 | pNH604-*pGPD1-LexA-B112* |
| pKW4019 | pFA6a-3xmKate2-*CaURA3MX6* |
| pKW4073 | pNH603-*pGPD1-LexA-B112* |
| pKW4190 | pRS313-*HR1_Chr2(SCO2)-p4xLexOcyc1-3xGST-V5-24xPP7sl-tCYC1-NatNT2-HR2_Chr2(SCO2)* |
| pKW4325 | pFA6a-*IAA7-3V5-KanMX6* |
| pKW4326 | pRS425-*p4xLexOcyc1-CDC33-3V5* |
| pKW4327 | pRS425-*p4xLexOcyc1-CDC33* |
| pKW4328 | pRS425-*p4xLexOcyc1-cdc33<sup>ΔG</sup>* |
| pKW4329 | pRS425-*p4xLexOcyc1-cdc33<sup>Δcap</sup>* |
| pKW4330 | pRS425-*p4xLexOcyc1-cdc33<sup>ΔGΔcap</sup>* |
| pKW4331 | pRS425-*p4xLexOcyc1-cdc33<sup>ΔG</sup>-3V5* |
| pKW4332 | pRS425-*p4xLexOcyc1-cdc33<sup>Δcap</sup>-3V5* |
| pKW4333 | pRS425-*p4xLexOcyc1-cdc33<sup>ΔGΔcap</sup>-3V5* |

**Supplemental Table 4.**
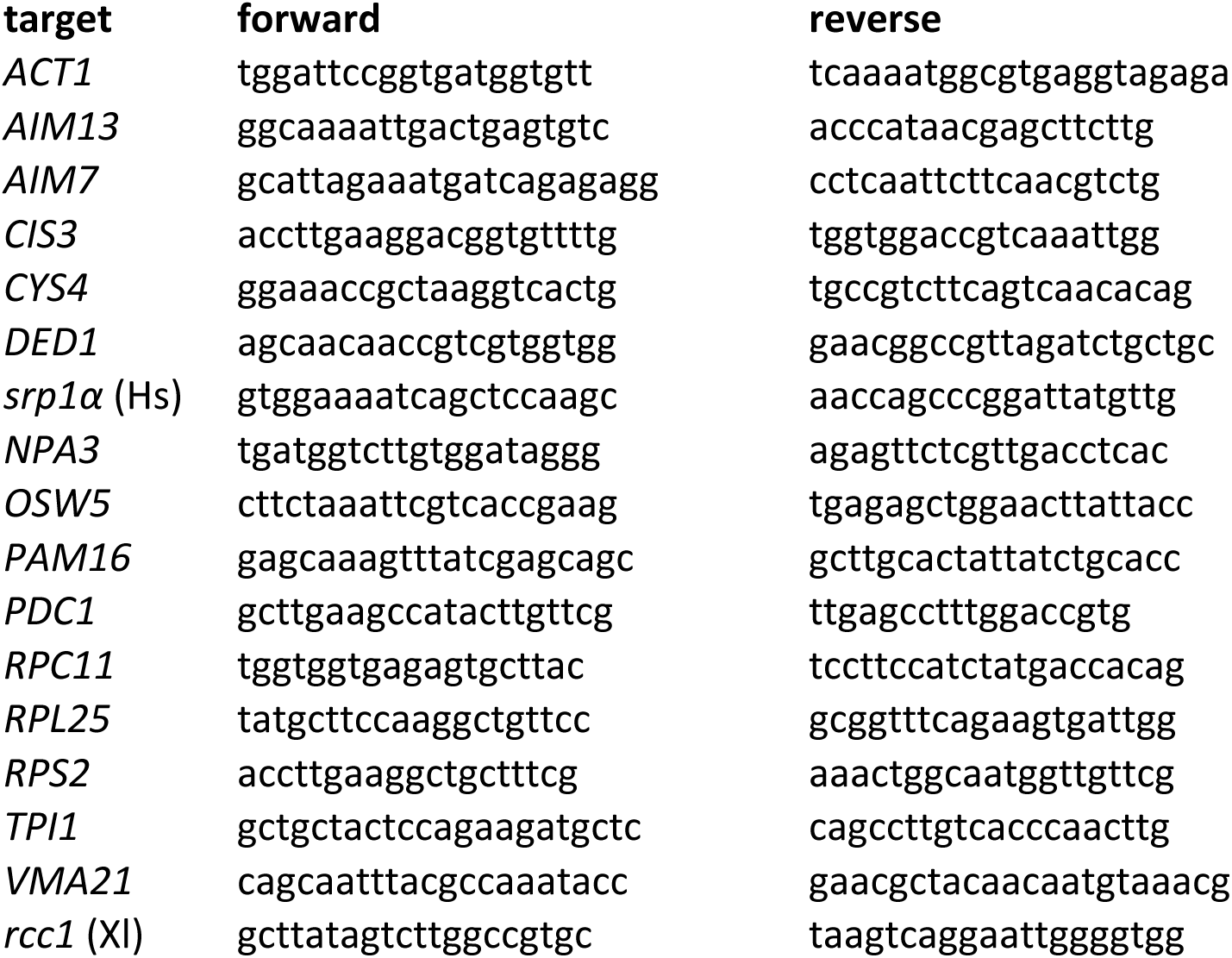

**Supplemental Figure 1.**
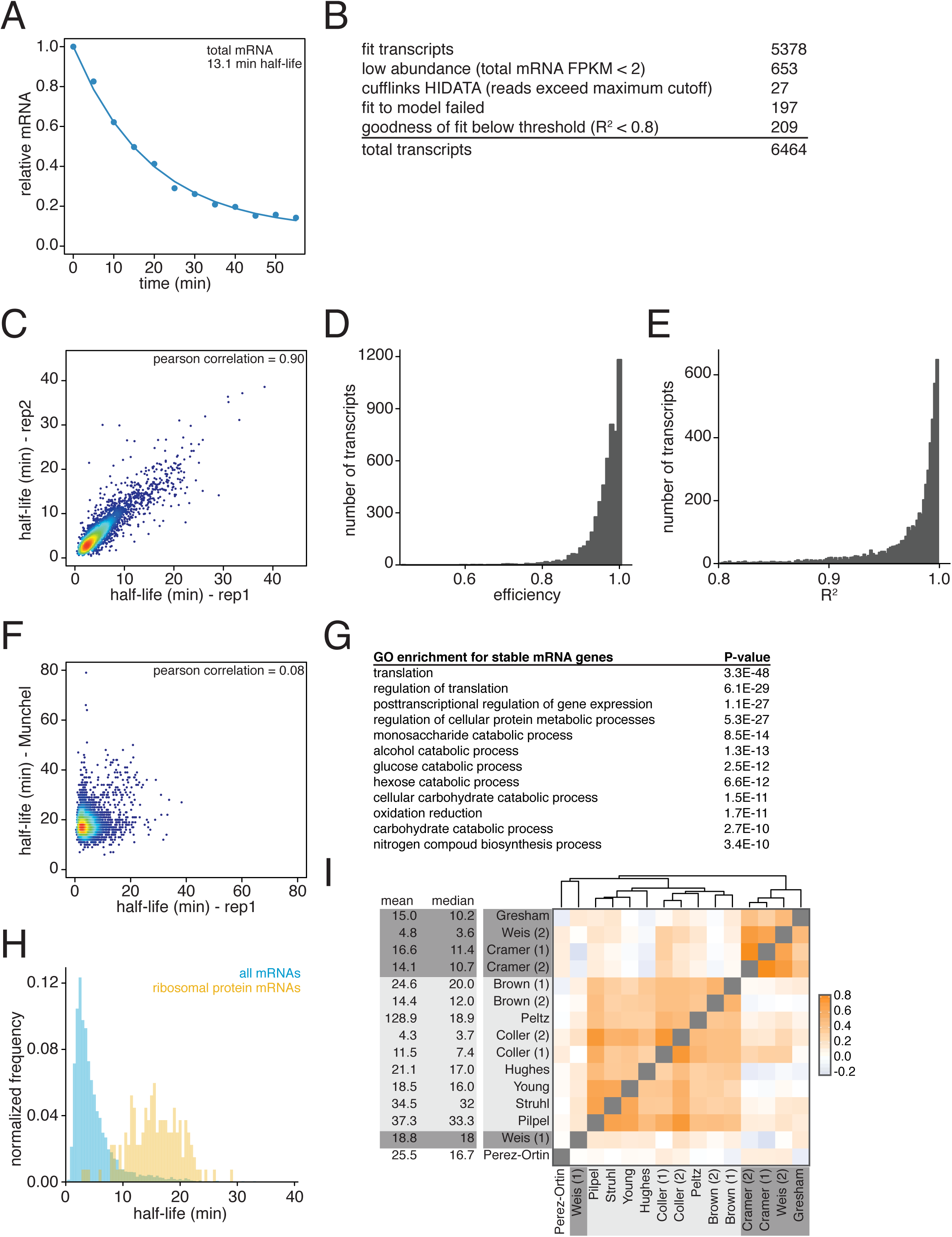
(A) Fragments per kilobase per million reads were summed for all mRNAs derived from the global stability profiling, normalized to the spike control and the time = 0 value, plotted and fit for half-life determination. (B) Numbers of transcripts that were successfully fit to the model and numbers of transcripts that failed to be measured for listed reasons. (C) Two biological replicates of the global mRNA stability profiling of wild-type cells (KWY165) were collected and transcript half-lives are plotted against each other. Red portions of the plot represent areas with dense datapoints and blue portions of the plot represent areas with sparse datapoints. (D) Distribution of efficiency parameters for the experiment described in Figure 1C. Each bin is 0.01 units wide. (E) Distribution of R^2^ values for the experiment described in Figure 1C. Each bin is 0.002 units wide. (F) Scatter plot of half-life values measured in this study compared to the values reported in Munchell et al 2011. Colors represent density of datapoints as in supplemental Figure 1C. (G) Functional enrichment of long-lived transcripts as defined as transcripts having a half-life longer than one standard deviation greater than the mean half-life. (H) Half-life distributions of all 137 ribosomal protein-encoding mRNA half-lives (yellow) compared to the entire transcriptome (blue) normalized to the size of each transcript group. Each bin is 0.5 minutes wide. (I) Spearman correlation coefficients were computed for pairs of mRNA stability datasets and plotted. Orange represents positive correlation and blue represents negative correlation. Text background colors represents experimental methodology: metabolic labeling in dark gray, transcriptional shutoff in light gray and other in white. The datasets were hierarchically clustered based on Euclidian distances. The datasets were derived from the following: Gresham [1], Weis (2) [this study], Cramer (1) [2], Cramer (2) [3], Brown (1) and (2) [4], Peltz [5], Coller (1) and (2) [6], Hughes [7], Young [8], Struhl [9], Pilpel [10], Weis (1) [11], Perez-Ortin [12]

**Supplemental Figure 2.**
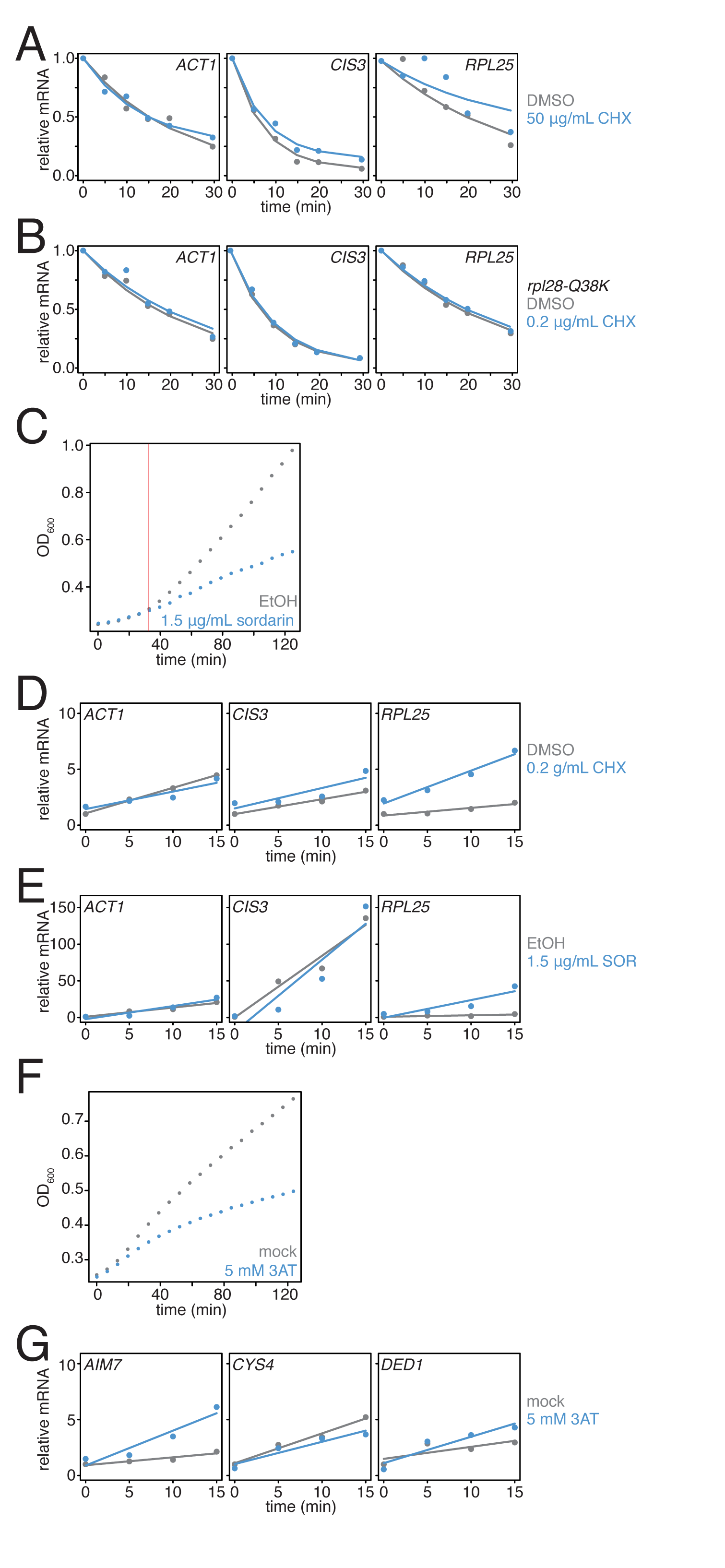
(A) Wild-type cells (KWY165) were subjected to mRNA stability profiling immediately after addition of 0.1% DMSO or 50 μg/mL cycloheximide in 0.1% DMSO. Data were collected and plotted as in Figure 3B. (B) *rpl28-Q38K* cells (KWY7335) were subjected to mRNA stability profiling immediately after addition of 0.1% DMSO or 0.2 μg/mL cycloheximide in 0.1% DMSO. Data were collected and plotted as in Figure 3B. (C) Wild-type cells (KWY165) were treated with 0.1% ethanol (gray) or 1.5 μg/mL sordarin in 0.1% ethanol (blue) and growth by absorbance at 600 nm was monitored. The horizontal orange line marks t = 33 min where the growth rates diverge. (D) Thio-uracil containing mRNAs from the experiment described in Figure 3A were measured and levels were plotted. RNA levels were normalized to a 4TU-labeled hSRP1 spike RNA and to the time = 0 value for the mock (DMSO) treated sample. (E) Thio-uracil containing mRNAs from the experiment described in Figure 3B were measured and levels were plotted. RNA levels were normalized to a 4TU-labeled hSRP1 spike RNA and to the time = 0 value for the mock (EtOH) treated sample. (F) Wild-type cells (KWY165) were treated with 5 mM 3AT (blue) or untreated (gray) and growth by absorbance at 600 nm was monitored. (G) Thio-uracil containing *AIM7*, *CYS4* and *DED1* mRNAs from the experiment described in Figure 3C-E were measured and levels were plotted. RNA levels were normalized to a 4TU-labeled hSRP1 spike RNA and to the time = 0 value for the mock (DMSO) treated sample.

**Supplemental Figure 3.**
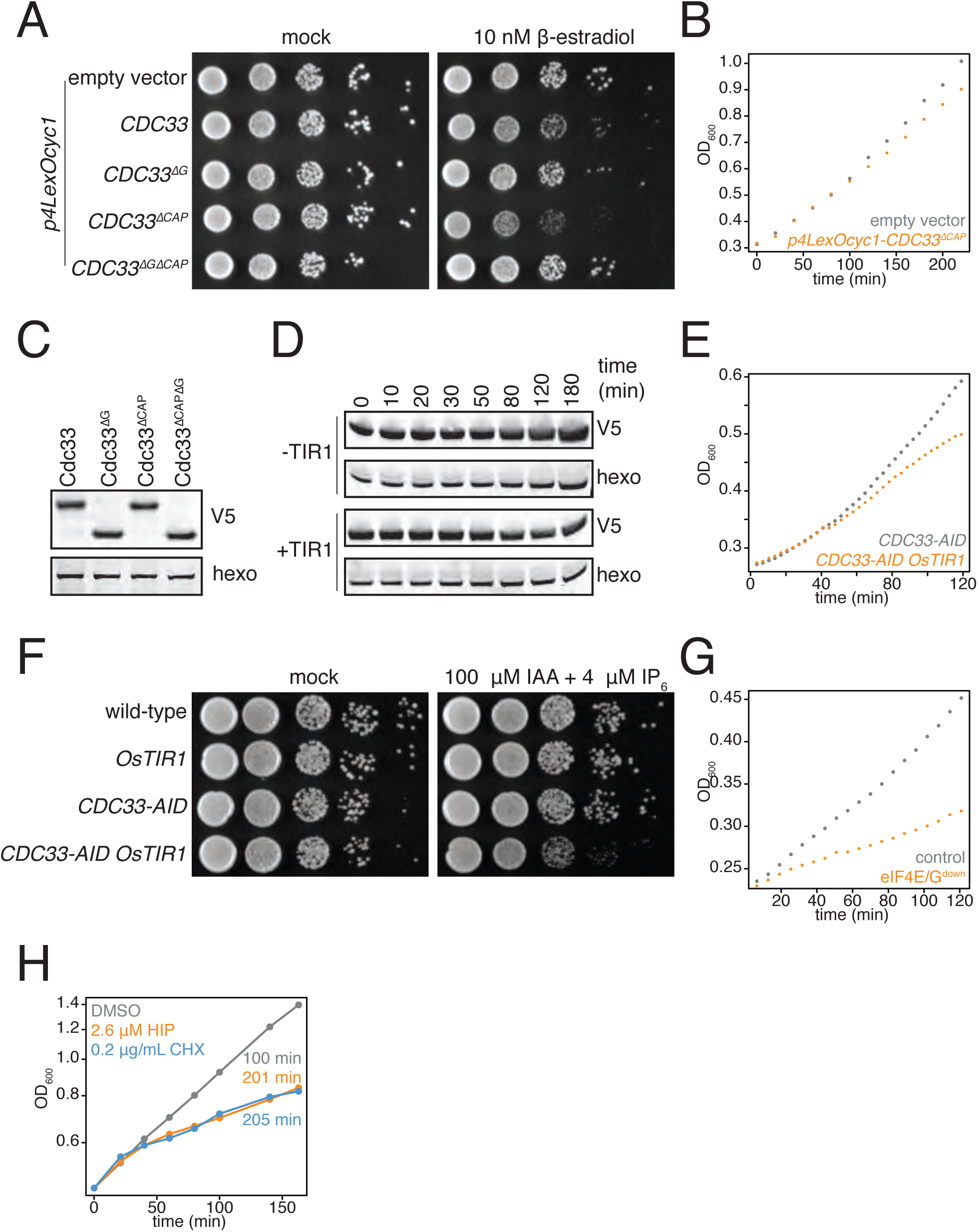
(A) *pGPD1-LexA-B112* cells harboring the following vectors: pRS425 (KWY7327 empty vector), pRS425-*p4LexOCYC1-CDC33* (KWY7329), pRS425-*p4LexOCYC1-CDC33*^*∆N*^ (KWY7330), pRS425-*p4LexOCYC1-CDC33*^*∆CAP*^ (KWY7331) and pRS425-*p4LexOCYC1-CDC33*^*∆CAP∆N*^ (KWY7332) were spotted on CSM-LEU and CSM-LEU + 10 nM β-estradiol media and incubated at 30°C for 36 hours. The first spot represents growth of approximately 6×10^4^ cells and each subsequent spot is a 10 fold serial dilution. (B) *pGDP1-LexA-B112* cells harboring pRS425 (KWY7327 empty vector, gray) or pRS425-*p4LexOCYC1-CDC33*^*∆CAP*^ (KWY7331, orange) were grown to mid exponential phase, treated with 10 nM β-estradiol and growth was monitored by absorbance at 600 nm. (C) pGDP1*-LexA-B112* cells harboring the following vectors: pRS425-*p4LexOCYC1-CDC33-3V5* (KWY7328), pRS425-*p4LexOCYC1-CDC33^∆N^-3V5* (KWY7338), pRS425-*p4LexOCYC1-CDC33^∆CAP^-3V5* (KWY7339) and pRS425-*p4LexOCYC1-CDC33^∆CAP∆N^-3V5* (KWY7340) were grown to mid exponential phase in CSM-LEU media, treated with 10 nM β-estradiol for 20 minutes and then harvested for western blot analysis. Hexokinase was used as a loading control. (D) *CDC33-IAA7-3V5* cells expressing OsTIR1 (KWY7326) or not expressing OsTIR1 (KWY7325) were grown to mid exponential phase in CSM pH5.5 media, treated with 100 μM 3-indoleacetic acid and 4 μM IP6 and then harvested at the indicated timepoints for western blot analysis. Hexokinase was used as a loading control. (E) *CDC33-IAA7-3V5* cells expressing OsTIR1 (KWY7326, orange) or not expressing OsTIR1 (KWY7325, gray) were grown to mid exponential phase in CSM pH5.5 media, treated with 100 μM 3-indoleacetic acid and 4 μM IP6 and growth by absorbance at 600 nm was monitored. (F) Wild-type (KWY165), *pGPD1-OsTIR1* (KWY7333), *CDC33-IAA7-3V5*(KWY7325) and *pGPD1-OsTIR1 CDC33-IAA7-3V5* (KWY7326) cells were spotted on YePD pH5.5 and YePD pH5.5 + 100 μM 3-indoleacetic acid and 4 μM IP6 media and incubated at 30°C for 36 hours. The first spot represents growth of approximately 6×10^4^ cells and each subsequent spot is a 10 fold serial dilution. (G) *pGPD1-LexA-B112 CDC33-IAA7-3V5* pRS425 (KWY7336 “control”, gray) and *pGPD1-LexA-B112 pGPD1-OsTIR1 CDC33-IAA7-3V5* pRS425-*p4xLexOcyc1-CDC33*^*∆CAP*^ (KWY7334 “eIF4E/G^down^”, orange) cells were grown in CSM –LEU pH5.5 to mid exponential phase, treated with 10 nM β-estradiol, 100 μM 3-indoleacetic acid and 4 μM IP6 and growth by absorbance at 600 nm was monitored. (H) Wild-type cells (KWY165) were treated with 0.1% DMSO (gray) or 2.6 μM hippuristanol (orange) or 0.2 μg/mL cycloheximide (blue) and growth was monitored by absorbance at 600 nm. The y-axis is in log scale. The values are the computed doubling times.

**Supplemental Figure 4.**
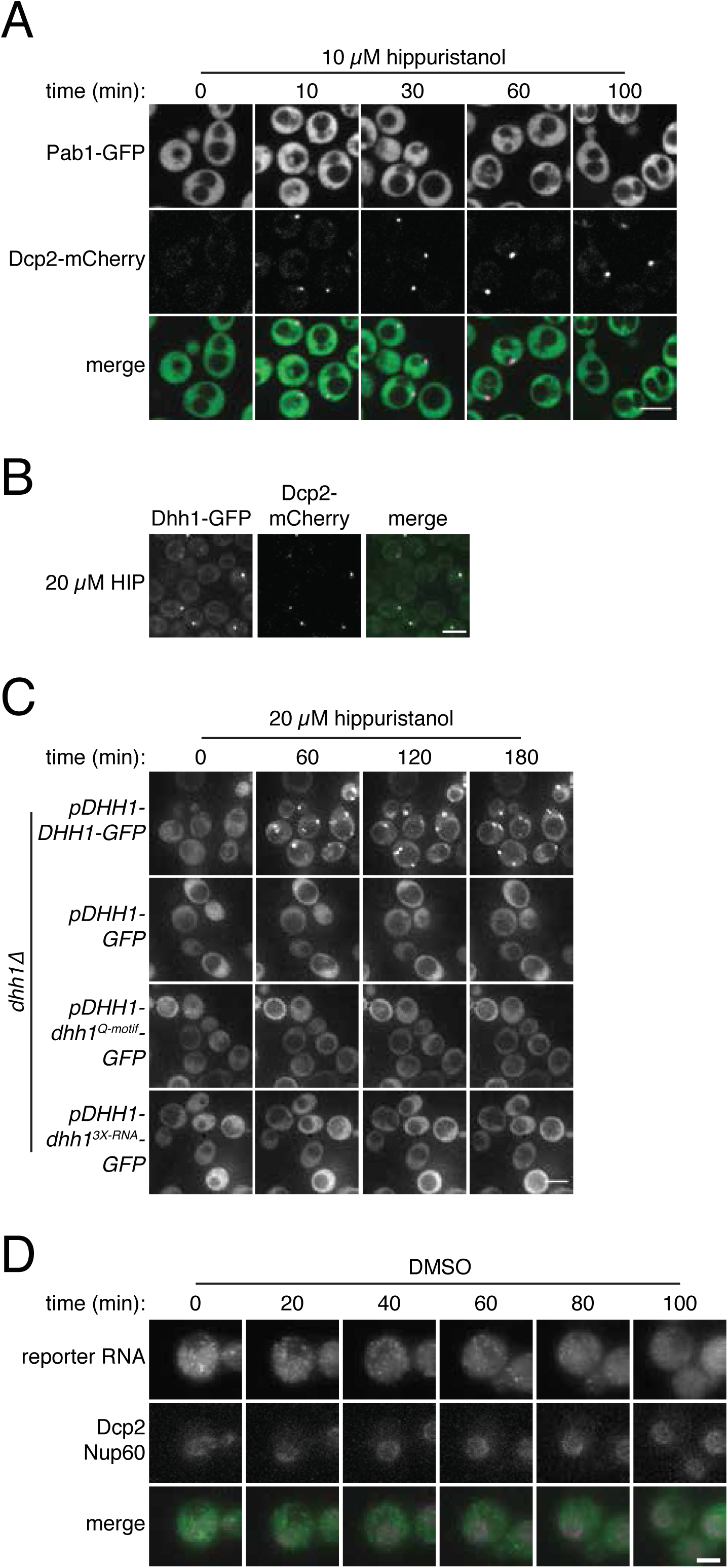
(A) Pab1-GFP, Dcp2-mCherry expressing cells (KWY6554) were grown to exponential phase, treated with 10 µM hippuristanol and imaged using a confocal microscope (n = 2 biological replicates). Scale bar: 5 μm. (B) Dhh1-GFP, Dcp2-mCherry expressing cells (KWY5948) were grown to exponential phase and then treated with 20 μM hippuristanol for 120 minutes. Imaging and image processing was performed as in Figure 4A (n = 4 biological replicates) Scale bar: 5 μm. Note that this image is from the same experiment depicted in Figure 4A. (C) *dhh1∆* cells expressing Dcp2-mCherry carrying plasmids *pDHH1-DHH1-GFP* (KWY3238), *pDHH1-GFP* (KWY5246), *pDHH1-dhh1^Q-motif^-GFP* (KWY5244) or *pDHH1-dhh1^3X-RNA^-GFP* (KWY5242) were grown to exponential phase, treated with 20 μM hippuristanol and samples were imaged at the indicated times. Imaging and image processing was performed as in Figure 4A (n = 3 biological replicates). Scale bar: 5 μm. (D) Dcp2-mCherry, Nup60-3xmKate2, PP7CP-GFP expressing cells carrying a synthetic 3xGST-24xPP7sl under β-estradiol inducible control (KWY7227) were grown to mid-exponential phase, treated with 400 nM β-estradiol for 40 minutes and then transferred to media lacking β-estradiol and containing 0.4% DMSO and immediately imaged. Imaging and image processing was performed as in Figure 4E (n = 4 biological replicates). Scale bar: 2 μm.

**Supplemental Figure 5.**
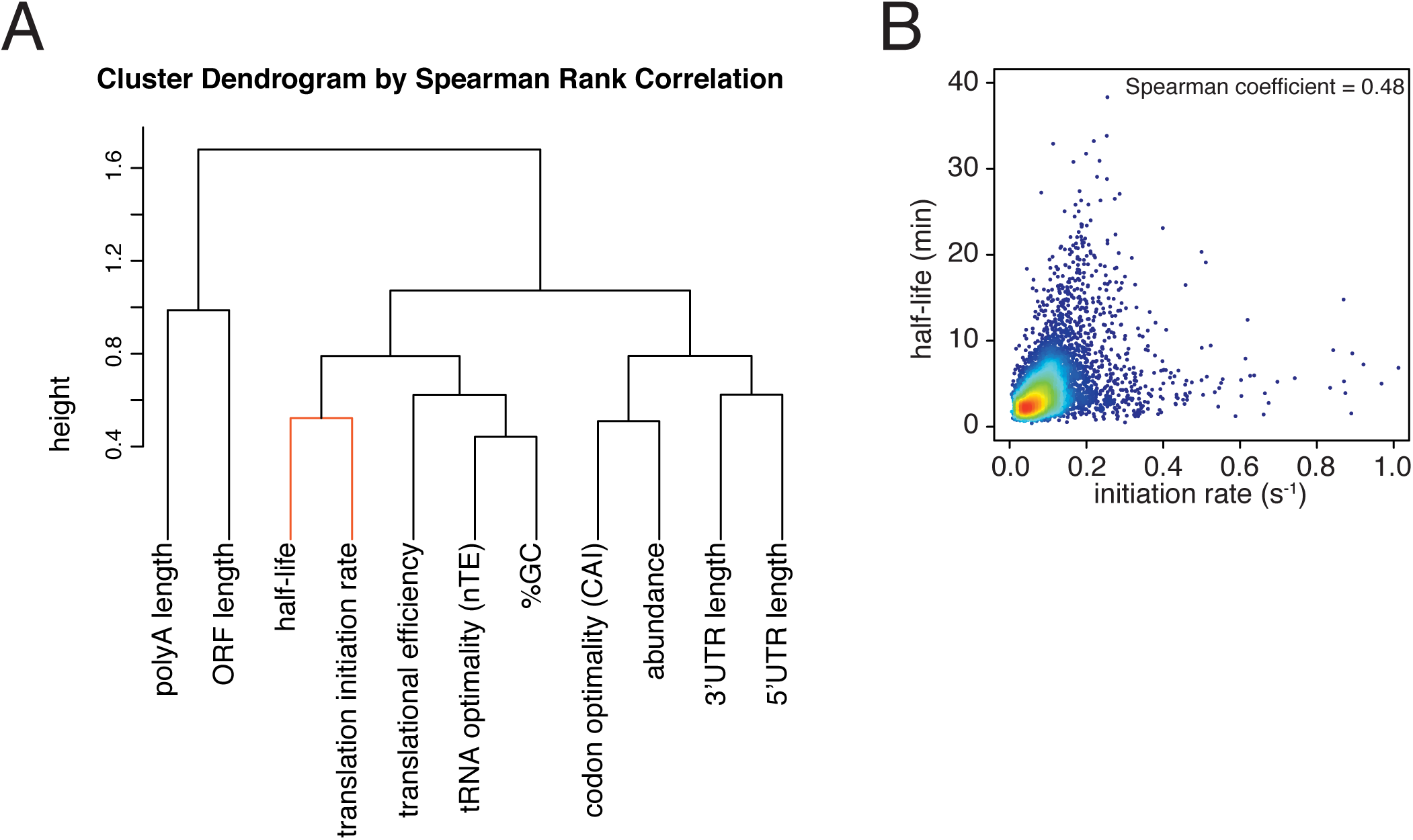
(A) Dendrogram of the Spearman rank correlation coefficients between transcriptome-wide datasets. Clustering was based on Euclidian distances. The close clustering between half-life and translation initiation rate is hilighted in orange. (B) Scatter plot of half-life values against estimated translation initiation rates. Color indicates density of datapoints as in supplemental Figure 1C.

## Extended technical supplement detailing the development and refinement of the 4TU-chase labeling experiment

To avoid the multiple problems associated with measuring mRNA stability using global transcriptional inhibition, we sought to employ metabolic labeling methods that would allow us to measure these kinetic parameters in a minimally invasive manner. We developed a working method and used this to collect one of the first profiles of mRNA stability in yeast cells [1]. Briefly, we grew cells to early exponential phase, labeled them with 0.2 mM 4TU for 2 hours and then transferred the cells to fresh media containing 19.6 mM (2.2 mg/mL) uracil. We collected cells from a timecourse during the chase phase and extracted total cellular RNA. We biotinylated these RNAs with HPDP-biotin, selected for the poly-adenylated fraction using oligo-dT beads and finally separated labeled mRNAs (here the “old-mRNA” pool) from unlabeled mRNAs (“newly-synthesized” pool). Based on follow-up work as well as the lack of agreement between our dataset and other metabolic-labeling based mRNA stability profiles, we sought to reexamine and improve our first-generation method.

To begin to test our method, we made use of the osmotic shock responsive gene, *STL1*. Upon osmotic shock, *STL1* mRNA is rapidly and transiently induced (Figure 1A and B) [2]. We performed our 4TU pulse-chase labeling and then subjected cells to an osmotic shock with the addition of 0.4M NaCl. If the pulse-chase labeling were performing optimally, we would expect that all *STL1* mRNAs would be contain only unmodified uracil. Rather, we found that a sizable fraction (about 40%) of *STL1* transcripts were eluted from the streptavidin beads (figure 2). This bleed through of a newly-made transcript into the labeled “old-transcript” pool could be a reflection of a number of possible technical problems. First, this could indicate that the chase phase is inefficient in preventing newly made transcripts from being labeled with 4TU. Second, this could indicate that the conjugation of biotin to the thiolated mRNA could be inefficient. Third, this bleed through could also be a symptom of carryover during the separation of thiolated mRNA from unlabeled mRNA. And fourth, it could be the case that our mathematical model requires modification to account for bleed through which cannot be addressed by experimental optimization.

The non-linear nature of detection based on affinity capture is inherent to the 4TU method and is a possible contributor to the bleed through problem. Put another way, in theory, an mRNA containing one thio-uracil could be treated the same as an mRNA containing 100 thio-uracils in the context of affinity purification. This is not an issue if the pulse-chase is performing optimally but even a small inefficiency in the chase, specifically new transcripts continuing to incorporate a low level of 4TU, would result in bleed through. We tested this possibility by again making use of the *STL1* system and varied the time of salt-shock and observed the degree to which newly made *STL1* mRNA partitioned between the bead-captured eluate and flowthrough fractions. We found that immediately upon uracil chase, salt shock led to a majority of newly made *STL1* transcripts being retained on the streptavidin beads. As we increased the lag time between uracil chase and salt-shock, we found that an increasing fraction of *STL1* mRNAs partitioned into the flowthrough (Figure 3A and B). The trend of increasing chase efficiency as time after chase was increased gave us a clear indication that inefficient chase was indeed a problem. We sought to correct this problem by switching from the existing 4TU-pulse/uracil-chase to a 4TU-chase labeling scheme where the non-linear nature of affinity capture-based detection is an advantage rather than a disadvantage. We grew cells to midexponential phase (OD600 = 0.4) and then added 0.2 mM 4TU to the culture. We again varied the time of salt-shock post 4TU chase and observed how the newly synthesized *STL1* transcript partitioned in the bead capture. We observe that even with no lag phase between 4TU chase and salt-shock, most of the newly synthesized *STL1* mRNA was effectively captured by the streptavidin beads. As the lag time was increased, a greater fraction of newly made *STL1* transcript was captured by the streptavidin beads (Figure 3C and D).

In addition to the switch in labeling scheme, we also tested the possibility that higher concentrations of 4TU in the chase could lead to improved new-transcript labeling. We titrated 4TU concentrations and found that the maximum tolerated dose of 4TU was 1 mM in our CSM-lowURA media (figure 4). Higher concentrations inhibited cell growth and would be self-defeating with respect to measuring mRNA dynamics in a minimally-perturbed system. We measured the stability of the *ACT1* transcript using a 0.2 mM 4TU-chase or a 1 mM 4TU-chase. Use of the higher 4TU concentration enabled a more efficient chase as indicated by the lower levels of *ACT1* transcript in the flowthrough fractions especially later in the timecourse (Figure 5).

We also examined if introducing a lag period between 4TU-chase and beginning the collection of the mRNA stability timecourse improved the efficiency of method. We measured the stability of the *ACT1* mRNA with a range of lag times. We found that even a brief lag period (2 minutes) improved the efficiency and consistency of the method (Figure 6). Increasing lag times beyond 2 minutes did not improve the quality of the data. Moreover, increasing lag times is a balancing act; some lag is beneficial for robust labeling but an overly long lag period fails to capture the dynamics of the most rapidly decayed mRNAs. Thus we opted for the 2 minute lag time where we still observed a beneficial effect.

Having optimized the labeling conditions, we next turned our attention to potential issues with carryover during the streptavidin bead-based RNA separation. We examined how unlabeled transcripts partitioned between the bead-bound and flowthrough fractions. We found that indeed unlabeled RNAs were being retained on the beads and moreover, different RNAs were retained to different degrees (compare Figure 3D lanes 1-3 for *STL1* and figure F7 lanes 1-3 for *rcc1 (Xl)*). This is especially problematic as it implies that in addition to a global distortion of mRNA stability, bead carryover can produce transcript specific artifacts as well thus complicating even relative comparisons of mRNA halflives. We reasoned that improvements to both bead blocking as well as bead washing could mitigate this carryover problem. We tested a range of blocking agents such as single-stranded salmon sperm DNA, polyA RNA, heparin and Denhardt’s reagent in addition to bacterial tRNAs as was used in our first-generation method. We found that bead-blocking with Denhardt’s reagent resulted in a marked improvement in unlabeled RNA carryover. Moreover, we found that this bead-blocking method did not interfere with the capture of an in vitro generated thio-labeled *srp1α (Hs)* RNA (Figure 7). We also examined how unlabeled RNAs were washed off the beads in the various steps. We found that most of the unlabeled RNA was contained in the flowthrough fraction. A fraction of remaining RNA that stuck to the beads was released with a high-salt 65C wash step but further washes with this buffer were ineffective in releasing additional non-specifically bound RNAs. We took advantage of the fact that the biotin-streptavidin interaction is resistant to SDS concentrations as high as 3%. With the addition of a single 1% SDS wash, we were able to release the majority of the remaining unlabeled RNA from the beads (Figure 8). Additional washes with this buffer did not release a significant amount of unlabeled RNAs. Again, we found that this more stringent wash protocol did not interfere with the ability of the streptavidin beads to properly capture labeled RNAs.

Recently, an improved biotin crosslinker, MTSEA-biotin, was developed and we tested if MTSEA-biotin performed better in our second-generation metabolic labeling method compared to the standard HPDP-biotin [3]. We collected a decay timecourse and processed the RNA in parallel only differing the biotin-crosslinker and analyzed the resulting decay curves. We found that the MTSEA-biotin resulted in better labeling evidenced by the reduced unlabeled mRNA levels we observed later in the timecourse (figures 9A and B).

Lastly, we revisited our computational model and examined the effects of what remaining inefficiencies in labeling and capture might have on simulated decay curves. We observed that as the efficiency of labeling and capture decreases, the curves become shallower and plateau at a non-zero steady state value (Figure 10A and C). Note that for a long lived transcript, fitting these data to a single exponential decay model can result in dramatically longer halflives with little loss of goodness of fit (Figure 10B). In the case of very unstable transcripts, the single exponential decay model completely fails for even modest levels of inefficiency (Figure 10D). To properly model these data, we introduced an efficiency of labeling and capture parameter to the standard decay model:

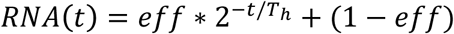

where *T*_*h*_ is the half-life and *eff* is a bulk efficiency of labeling and thio-RNA capture. Note that in the limiting case where the efficiency approaches 1, the equation reduces to the single exponential decay model. We used this two parameter model for all of our half-life determinations. Moreover, we did not use a global efficiency parameter but rather fit each transcript to its own efficiency as there are transcript specific contributions to this bulk parameter such as bead-stickiness and uracil content.

We made one final modification to the 4TU method with regard to spike-in control RNAs that normalize for differences in the various RNA recovery and bead binding steps of the 4TU protocol. In our first generation protocol, we spiked in control RNAs relative to a constant amount of total RNA which is comprised mostly of rRNA. We then accounted for the mRNA “decay by dilution” due to cell growth and division by correcting for the growth rate. This method introduces the errors of growth rate determination as well as RNA quantification into the protocol. We reasoned that in theory, adding our spike-in controls relative to a constant **culture volume**, would eliminate the need to correct for “decay by dilution.” The only caveat to this approach is the assumption that the efficiency of cell lysis is relatively constant. We set out to test this assumption. We found that for lysing 6 identical cell pellets that there was a 3% standard deviation in RNA recovery (Figure 11). We also tested the linearity of recovery of RNA from increasing sizes of cell pellets and found that that the recovery of RNA was robustly linear to input cell material over the range of our experiments (Figure 12). We concluded that spiking relative to culture volume is an improvement on the first-generation metabolic labeling protocol.

In total, we re-examined each step of our 4TU protocol and made improvements where possible. We then introduced a modified mathematical model to account for the remaining inefficiencies in the protocol that we could not experimentally correct. This has resulted in an improved second generation 4TU protocol that is accurate and reproducible.

## Experimental Procedures

### Northern blot analysis

Northern blot experiments were performed as previously described (Carroll et al. 2011). Oligonucleotide probes are as follows:

*STL1* GATTTTGGGACCTGCCTCTGGAGAACAAACTTGACAGTG

*rcc1 (Xl)* GAAAGACCAAAGCCATATACATGGCCTTCTTGGGACACCGC

*srp1* α (Hs) GGGCTGTTTTTCTCTGGAAAGTAGTTTCCTGGCAGCTTGAG

### Figure Legends

**Figure 1:**
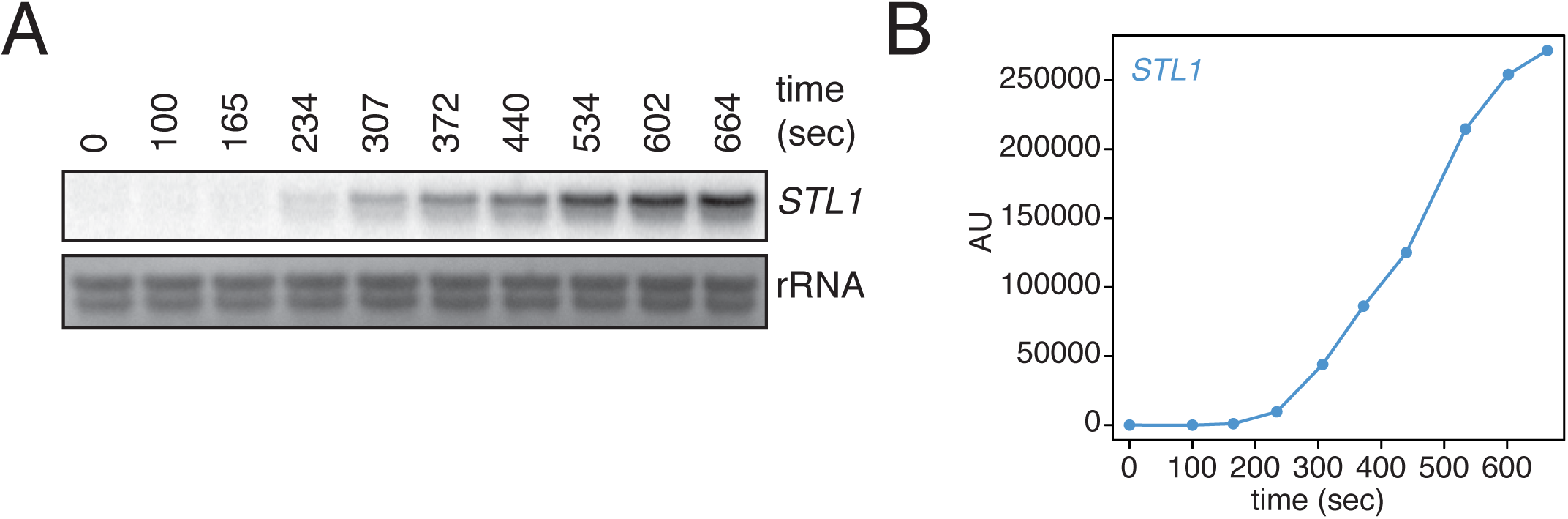
Kinetics of *STL1* mRNA induction. (A) Wild-type cells (KWY165) were grown to exponential phase, subjected to a 0.4 M NaCl salt shock and samples were collected at the indicated times for northern blot analysis. (A) Quantification of data in (A). *STL1* mRNA levels were corrected to background and also to rRNA levels.

**Figure 2:**
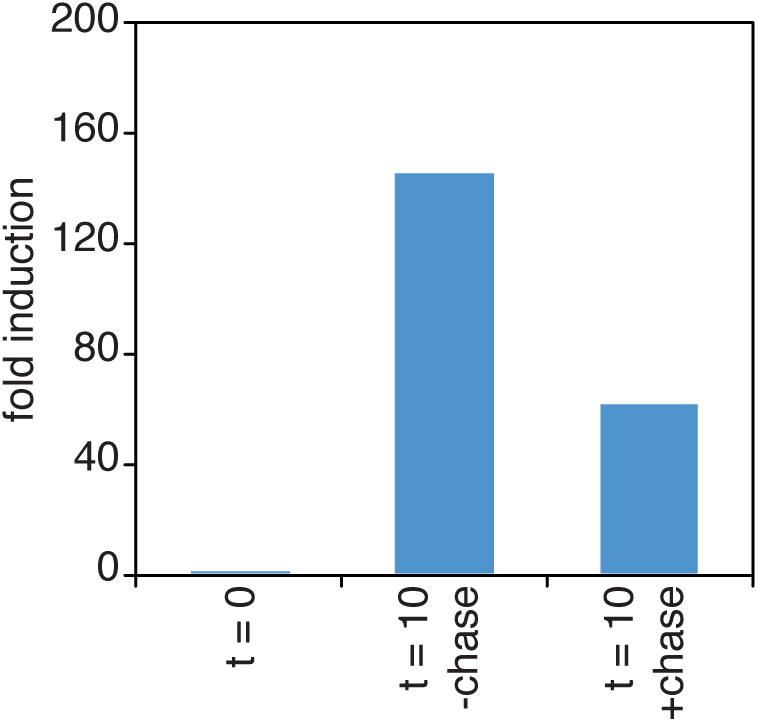
Chase inefficiency as revealed by *STL1* induction. Wild-type cells (KWY165) were grown to OD600 = 0.2, labeled with 0.2 mM 4TU for 2 hours and then washed out of the labeling media. Half of the cells were returned to the same media and the other half were resuspended in chase media containing 19.6 mM uracil. Both cultures were immediately subjected to 0.4 M NaCl salt shock and samples were collected 10 min after the salt shock. Thio-labeled mRNAs were purified and abundance of *STL1* thio-mRNA was determined by qPCR, normalized to a thio-labeled spike (*srp1α (Hs)*) and then normalized to the levels of *STL1* in pre-salt shocked cells (t = 0).

**Figure 3:**
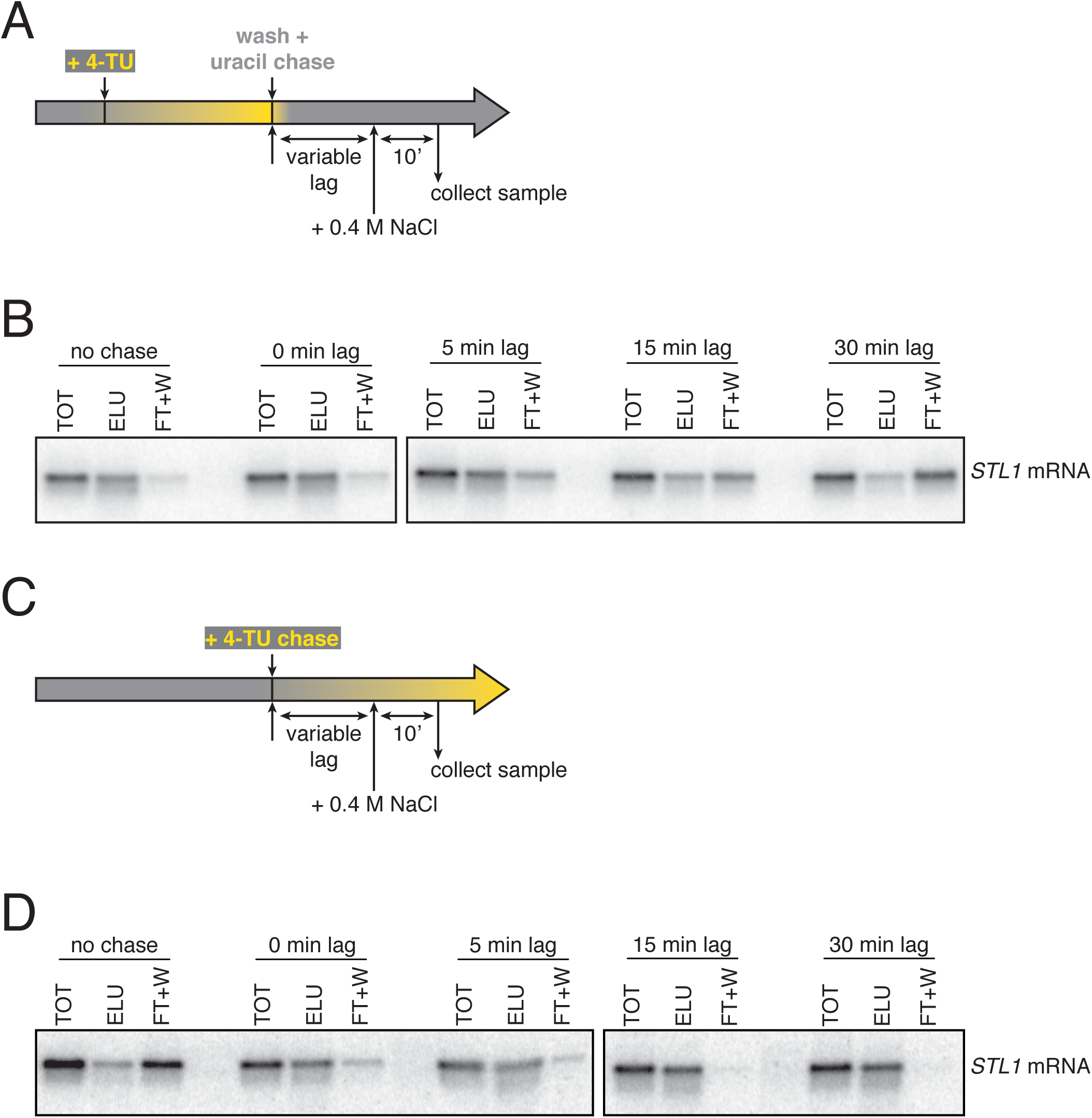
Analysis of chase efficiencies as a function of lag time and labeling scheme. (A) 4TU-pulse/uracil-chase labeling scheme and experimental setup for (B). (B) Wild-type cells (KWY165) were grown to exponential phase, the experiment outlined in (A) was performed and levels of *STL1* mRNA were determined by northern blot. (C) 4TU-chase labeling scheme and experimental setup for (D). (D) Wild-type cells (KWY165) were grown to exponential phase, the experiment outlined in (C) was performed and levels of *STL1* mRNA were determined by northern blot.

**Figure 4:**
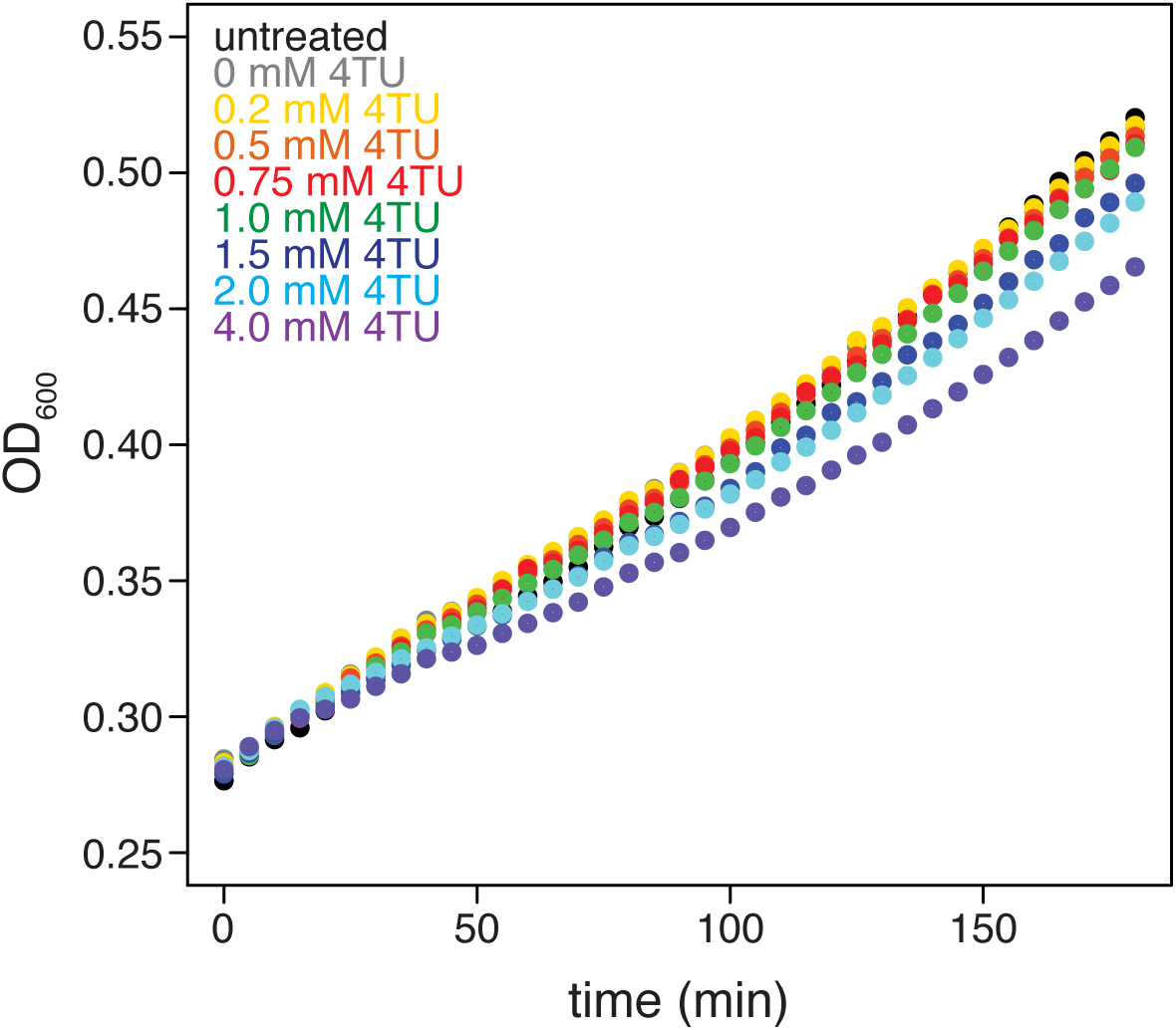
Effects of 4TU on cell growth. Wild-type cells (KWY165) were grown to exponential phase, treated with the indicate concentration of 4TU and growth was monitored by absorbance at 600 nm.

**Figure 5:**
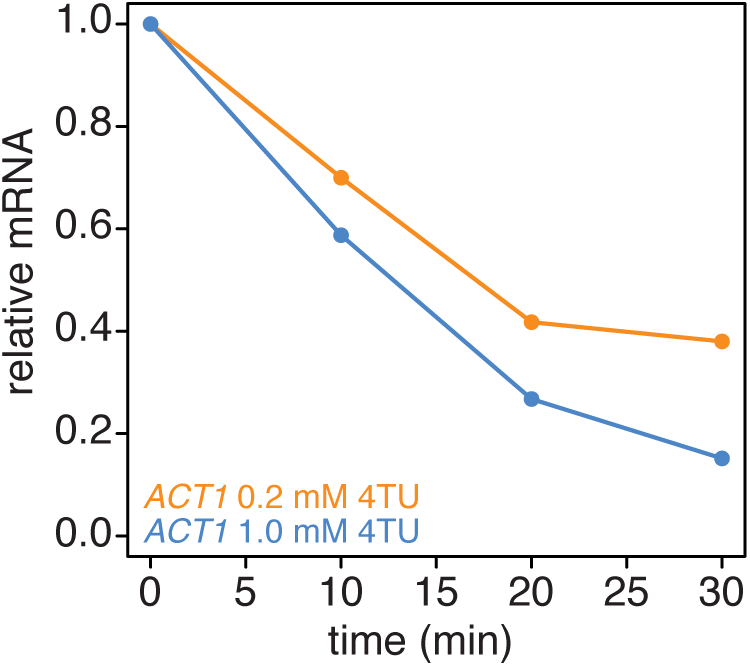
Increasing the 4TU concentration during the chase improves the subtraction of newly synthesized mRNAs. Wild-type cells (KWY165) were subjected the 4TU-chase protocol with either 0.2 mM 4TU in the chase (orange) or 1 mM 4TU (blue) and the unlabeled *ACT1* mRNA levels were quantified.

**Figure 6:**
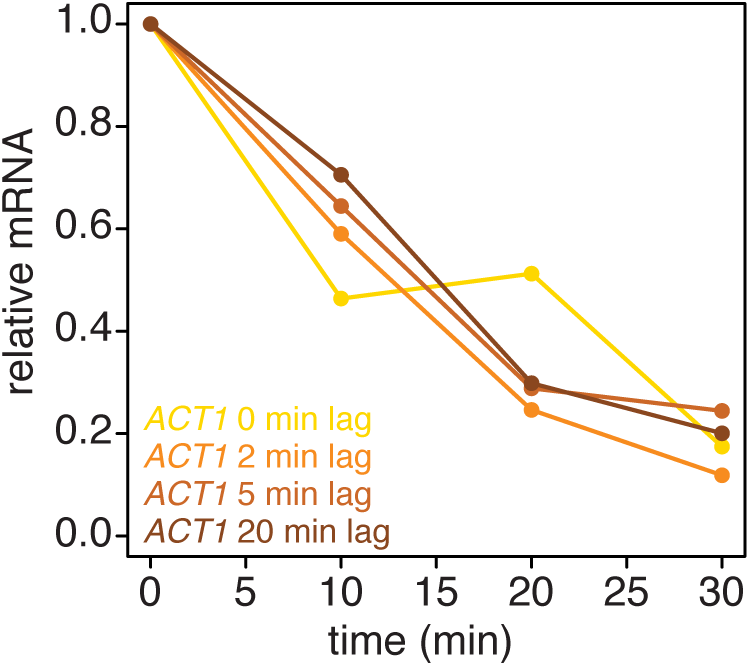
Introducing a brief lag period between the 4TU-chase and starting the decay timecourse improves the quality of the decay data. Wild-type cells (KWY165) were subjected the 4TU-chase protocol. A variable amount of time was allowed to elapse after the 4TU-chase and collection of the t = 0 sample and the unlabeled *ACT1* mRNA levels were quantified.

**Figure 7:**
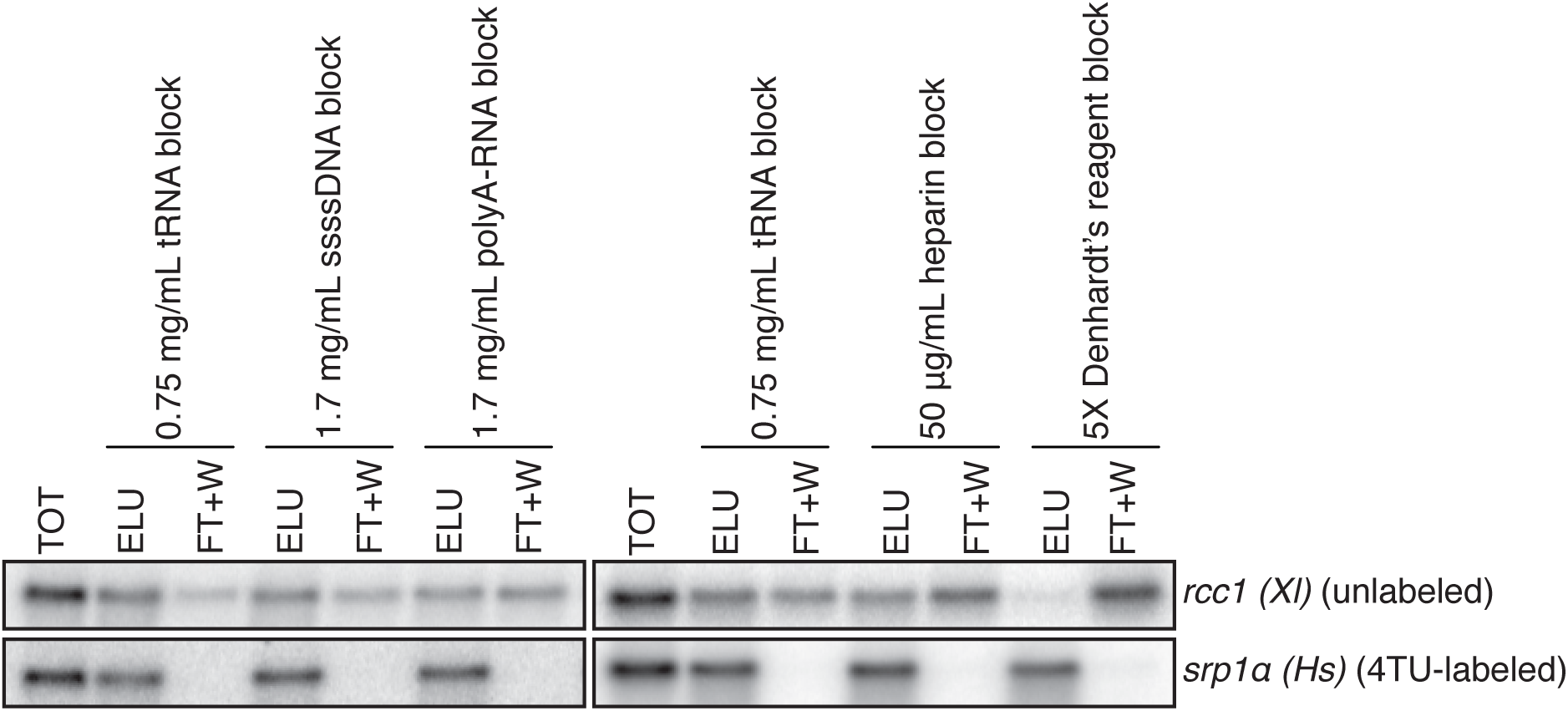
Analysis of streptavidin bead blocking efficiency. Wild-type cells (KWY165) were labeled with 1 mM 4TU for 2 hours. 5 ng of in vitro transcribed *rcc1 (Xl)* mRNA spike and 5 ng of in vitro transcribed 4TU-labeled *srp1α (Hs)* mRNA spike were added to 10 ug of extracted total RNA and the RNA mix was biotinylated. mRNAs were enriched for using oligo-dT beads and the mRNAs were then subjected to streptavidin bead selection. Streptavidin beads were prepared using the indicated blocking agents and the total (TOT), eluate (ELU) and flowthrough plus washes (FT+W) were analyzed by northern blot.

**Figure 8:**
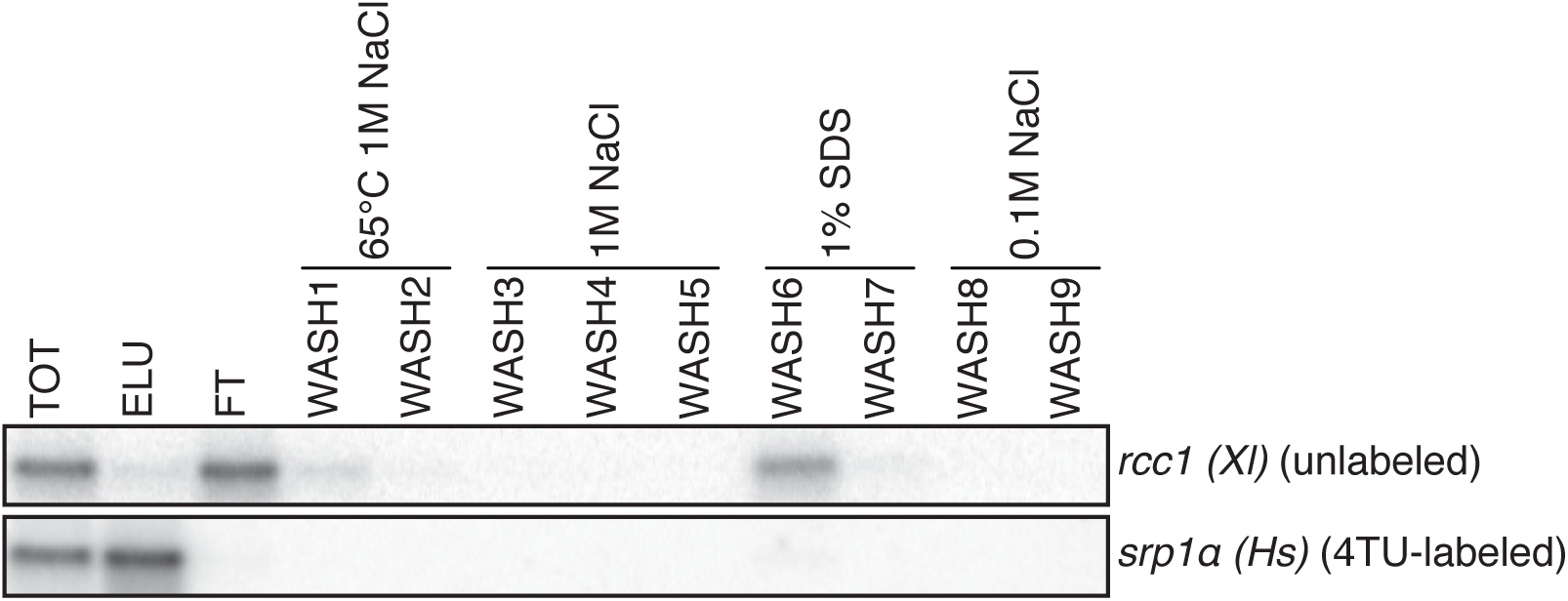
Analysis of unlabeled mRNA release during streptavidin bead purification. RNA mixtures were prepared as in Figure 7 and total (TOT), eluate (ELU) and washes (WASH1-WASH9) were analyzed by northern blot.

**Figure 9:**
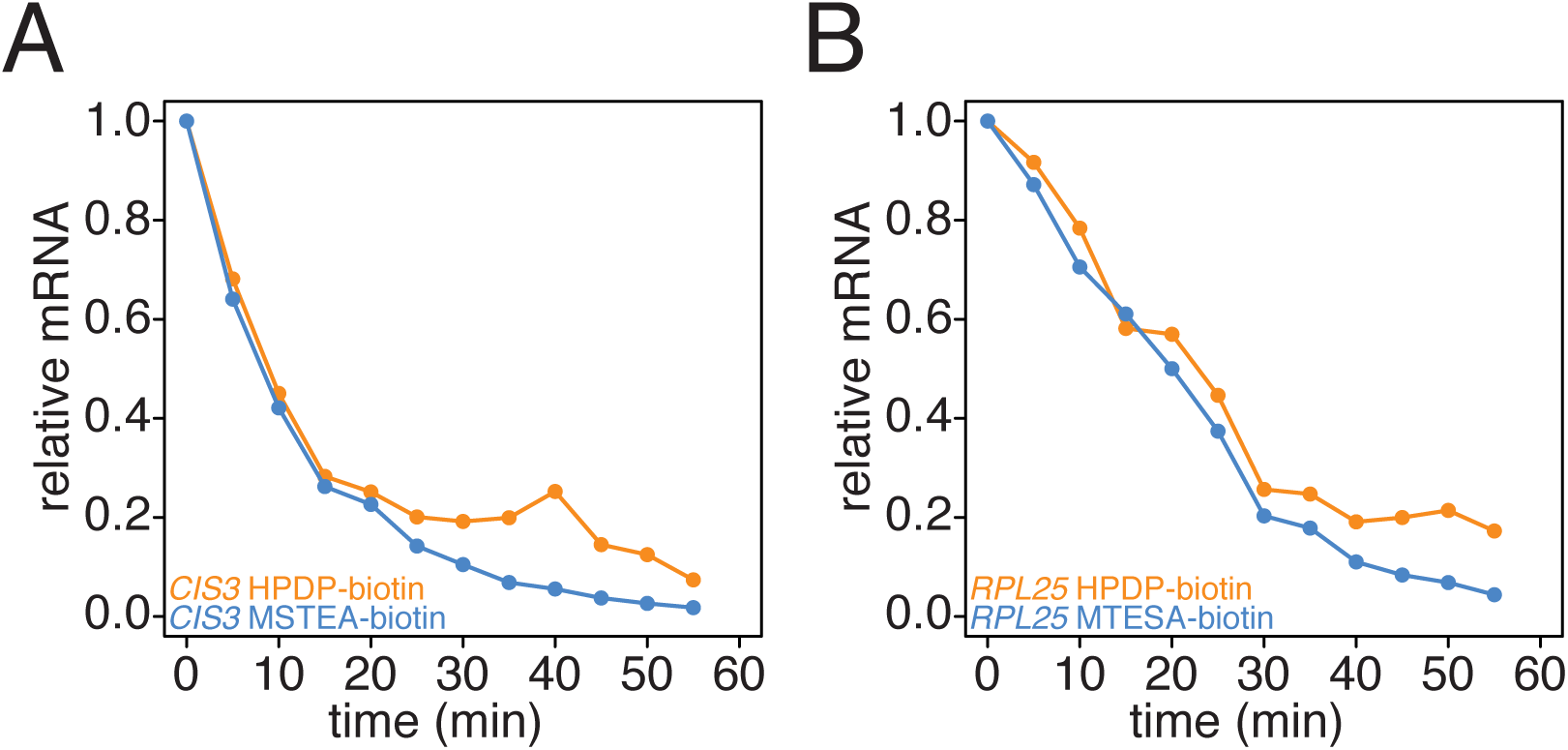
Comparison of HPDP-biotin with MTSEA-biotin in the efficiency of subtracting newly synthesized mRNAs during the chase phase. Wild-type cells (KWY165) were subjected the 4TU-chase protocol and RNA mixes were biotinylated with either HPDP-biotin (orange) or MTSEA-biotin (blue). Levels of unlabeled *CIS3* and *RPL25* mRNAs were determined.

**Figure 10:**
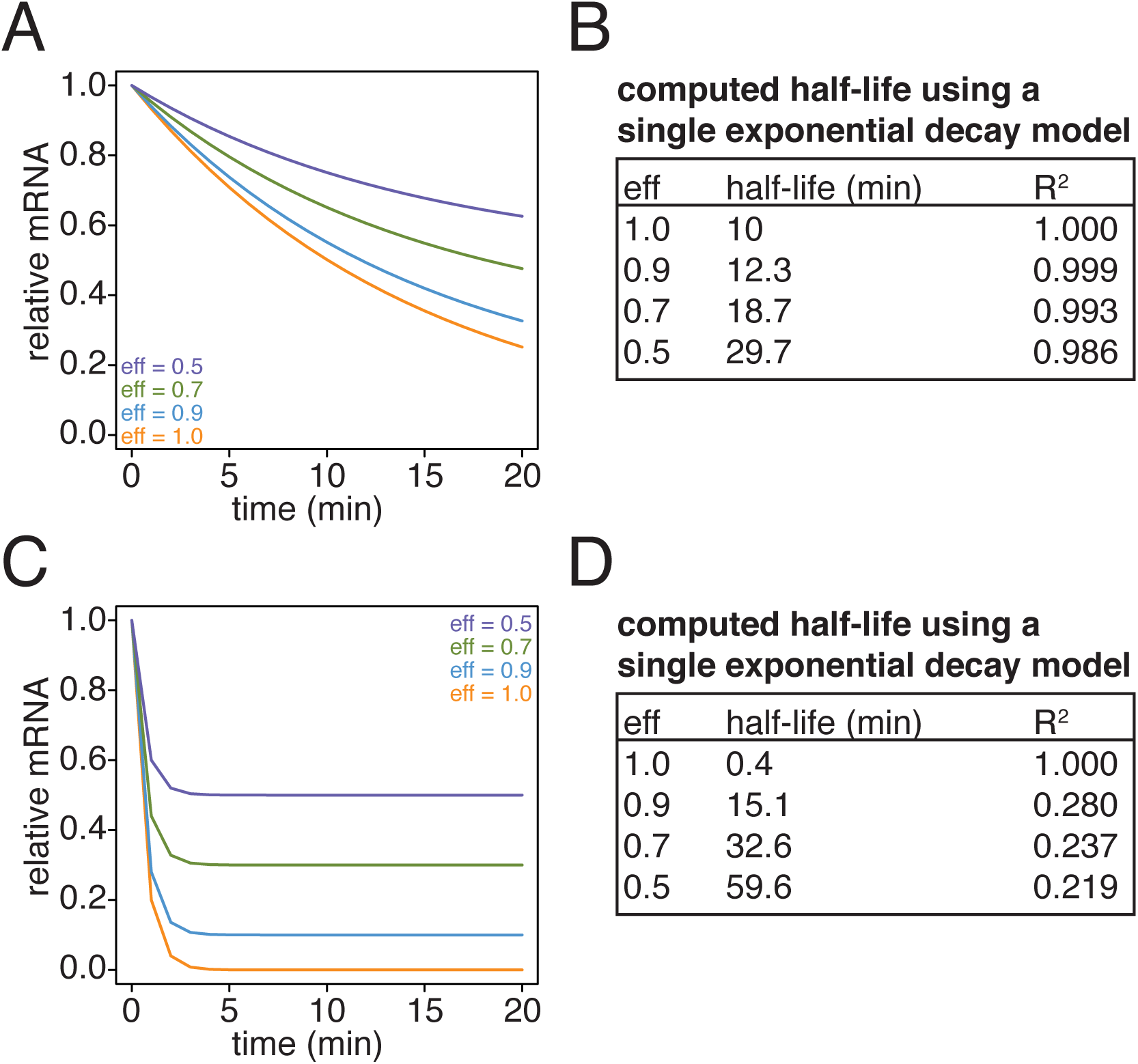
Simulation of the effects of non-ideal efficiencies in labeling and streptavidin bead separation on decay kinetics. (A) Decay data for an mRNA with a 10 minute half-life were simulated with variable degrees of labeling and purification efficiencies. (B) The simulated data in (A) were fit to a single exponential decay model (RNA(t) = RNA(0)*2^−t/hl^ where hl is the halflife) and the halflives and goodness of fits (R^2^) were determined. (C) Decay data for an mRNA with a 0.4 minute half-life were simulated with variable degrees of labeling and purification efficiencies. (D) The simulated data in (C) were fit to a single exponential decay model (RNA(t) = RNA(0)*2^−t/hl^ where hl is the halflife) and the halflives and goodness of fits (R^2^) were determined.

**Figure 11:**
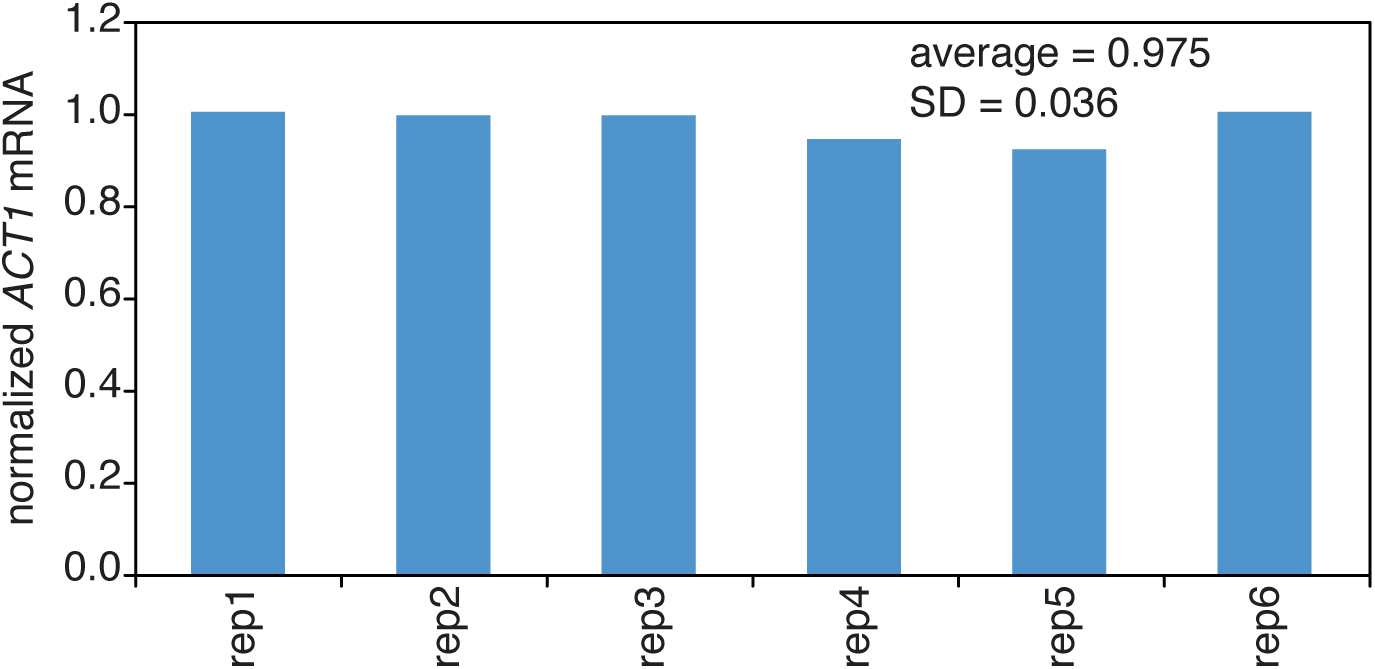
Reproducibility of cell lysis during RNA extraction. Wild-type cells (KWY165) were grown to exponential phase and 6 technical replicate cell pellets of 5 OD600 units were collected. 10 ng of in vitro transcribed *rcc1 (Xl)* mRNA spike was added to each sample and the cells were subjected to the RNA extraction protocol. Levels of *ACT1* and the spike mRNAs were determined and the ratio of *ACT1*:spike normalized to replicate 1 is plotted.

**Figure 12:**
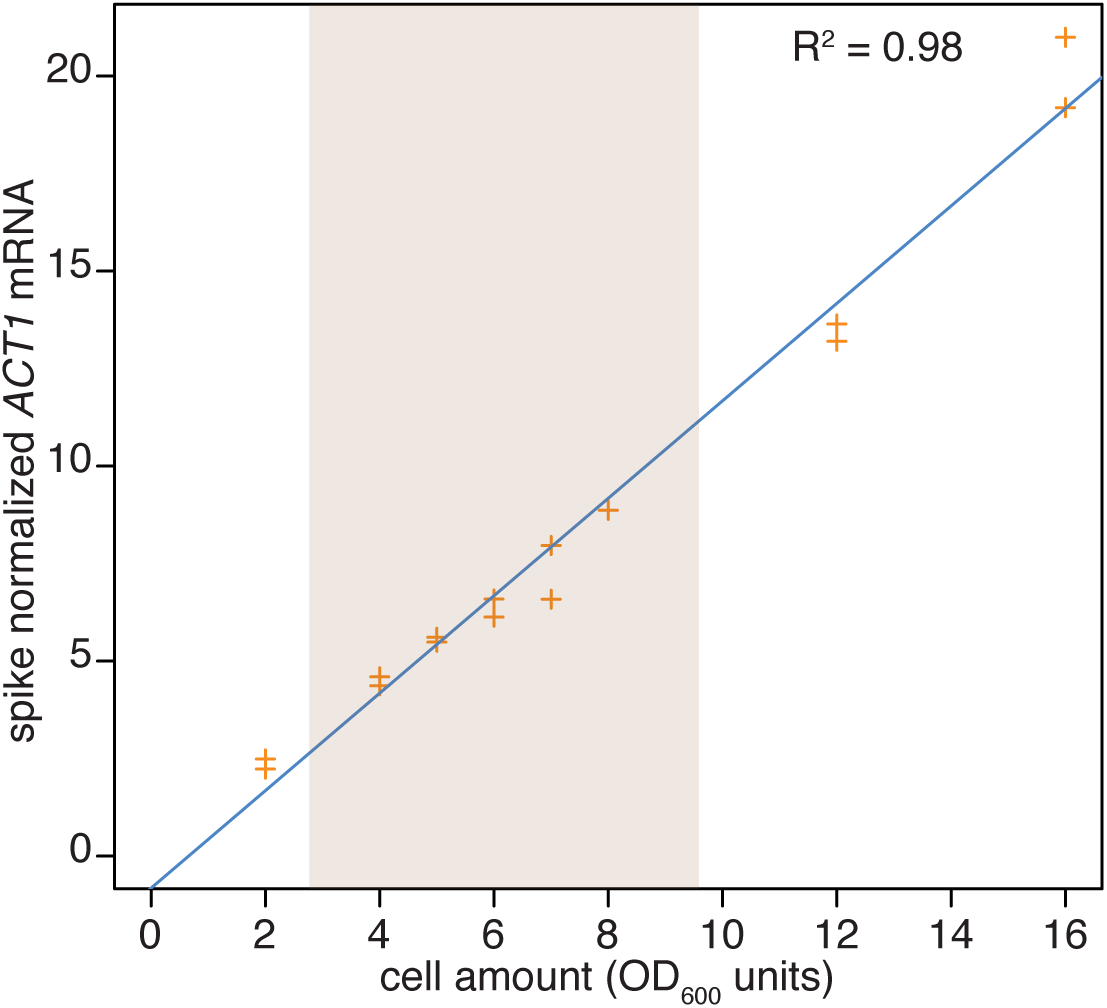
Reproducibility of cell lysis during RNA extraction. Wild-type cells (KWY165) were grown to exponential phase and 2 technical replicate cell pellets of varying OD600 units were collected. 10 ng of in vitro transcribed *rcc1 (Xl)* mRNA spike was added to each sample and the cells were subjected to the RNA extraction protocol. Levels of *ACT1* and the spike mRNAs were determined and the ratio of *ACT1*:spike is plotted. The shaded area indicates the OD600 range in which the 4TU-chase experiments are performed.

